# Self-supervised representations reveal the genetic architecture of human cortical folding

**DOI:** 10.64898/2026.07.23.740040

**Authors:** Antoine Dufournet, Julien Laval, Joël Chavas, Clara Fisher, Denis Rivière, Vincent Frouin, Jean-François Mangin

## Abstract

Cortical folding emerges during fetal development, is under genetic control, and remains stable throughout life, offering a lasting window into early neurodevelopment. Conventional morphometric descriptors, however, only partially capture the shape variability of cortical folds. Here, we compare 56 regional self-supervised deep learning representations of cortical folds with classical sulcal morphometry via multivariate genome-wide association studies (GWAS) in 35,940 UK Biobank participants. The learned representations identified 567 independent genome-wide significant loci, versus 162 for classical morphometry, 87% of which were also detected by our approach. More than half of these associations replicated in the independent Adolescent Brain Cognitive Development (ABCD) cohort. Gene, gene-set, BrainSpan and single-cell expression enrichment converge on a shared prenatal window of neurogenesis and morphogenesis; spatial gene-association maps recapitulate known regional expression gradients, including for *NR2F1*. Together, these results establish self-supervised representations of cortical folding as a powerful phenotypic framework for the genetic study of neurodevelopment.

## 2 Introduction

The human cerebral cortex exhibits a characteristic folded organisation, the emergence of which begins around the twentieth week of gestation [1–5]. Primary folds, which are the most conserved across individuals, delineate the major lobes and principal functional regions. Secondary and tertiary folds subsequently emerge between these primary folds and display increasing inter-individual variability [1, 4]. This variability is itself informative: monozygotic twins share markedly more similar folding patterns than dizygotic twins, demonstrating genetic control over the process [6–8]. Though the mechanisms underlying fold formation remain incompletely understood, they are thought to arise in part from regional fluctuations in progenitor cell proliferation, as well as neuronal migration and differentiation during fetal development. These processes might induce local growth differences that, under physical constraints imposed by tissue properties, would give rise to cortical folding [9, 10]. Furthermore, once established, cortical folding patterns remain remarkably stable throughout adulthood [11, 12]. Cortical folding is therefore a stable, genetically influenced trace of events that occurred almost exclusively during fetal life, making it a biomarker of early neurodevelopment that remains legible in an adult Magnetic Resonance Imaging (MRI) scan decades later.

Conventional morphometric descriptors of cortical anatomy do not fully exploit this substrate. Folds opening increases with age [13], and cortical thickness changes across the lifespan [14], while regional surface area summarises fold shapes into a single scalar per region. More broadly, handcrafted descriptors, such as surface area, volume, folds length, and depth, capture only a partial projection of the underlying folding patterns. A phenotype that is both stable across the lifespan and expressive enough to retain the local patterns of individual folds would provide a sharper, less noisy window into the genetic architecture of early neurodevelopment.

Among possible phenotypes, the cortical folding skeleton, a negative cast of the cortex [15–17], faithfully retains individual folding patterns (Fig. 1, step 1), without incorporating cortical thickness or folds opening. This phenotype can be automatically extracted from standard T1-weighted MRI scans acquired in both research and clinical settings, making it well suited for large-scale analyses in cohorts such as UKB [18] and ABCD study [19]. What it lacks, as a raw 3D object, is a compact and analysable representation. Representation learning offers it: deriving compact, data-driven phenotypes directly from high-dimensional observations, rather than relying on predefined morphometric measurements. Recent studies have shown that self-supervised methods can extract compact anatomical embeddings that improve downstream analyses across a range of imaging applications [20–22]. Whether such learned representations capture biologically meaningful genetic variation beyond conventional phenotypes, however, remains largely unexplored.

**Figure 1:**
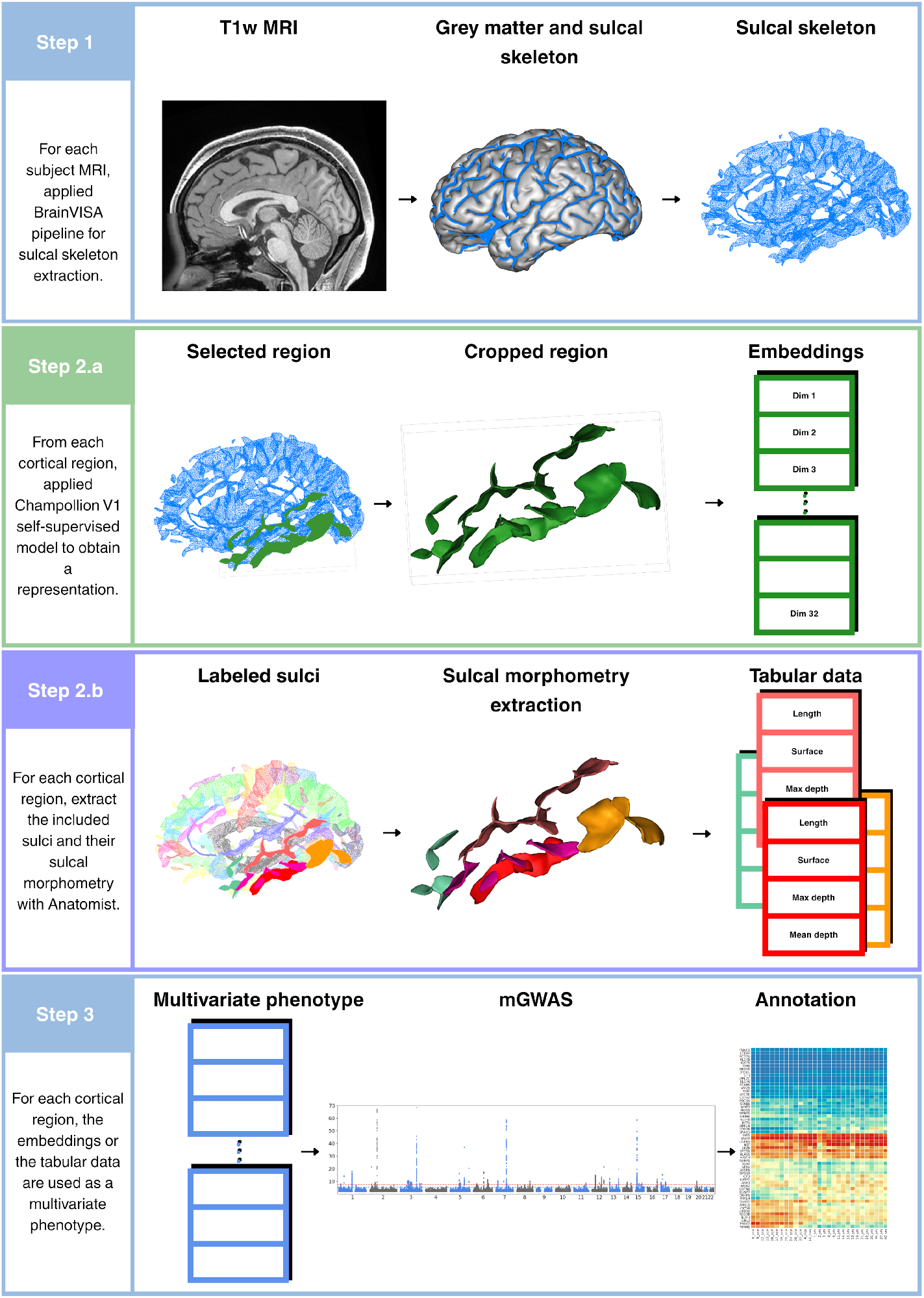
Overview of the analytical pipeline. Step 1: Extraction of cortical folding skeletons from T1-weighted MRI using the Morphologist v5.0/6.0 pipeline (BrainVISA). Each skeleton (in blue) is a binary volumetric representation of folding patterns in MNI space (2 mm isotropic resolution). Step 2a: Champollion V1 contrastive self-supervised framework encodes each of 56 cortical regions as a 32-dimensional representation. Step 2b: Classical sulcal morphometric descriptors (length, surface area, mean depth, maximum depth) are extracted per sulcus and aggregated per region. Step 3: Pre-residualization for covariates (age, age^2^, sex, age^2^ sex, TIV, imaging centre, first 20 genetic PCs), followed by a multivariate GWAS using MOSTest on the 32-dimensional representation space (2.a) or tabular data (2.b) of each region and downstream analyses including gene annotation (Multi-marker Analysis of GenoMic Annotation MAGMA), gene-set enrichment, BrainSpan expression enrichment, and genetic correlation analyses.

Here, we use Champollion V1 [8], a self-supervised deep learning framework trained on 42,000 UKB folding skeletons. It produces a 32-dimensional representation of cortical shape variation across the population (Fig. 1, step 2.a) for each of 56 cortical regions defined by an atlas of sulci and gyri. These representations show significant correlations with sulcal morphometric descriptors (Fig. 1, step 2.b), such as sulcal length, depth, and surface area [8], while also encoding shape and positional information that these descriptors discard. In contrast to approaches relying on strong non-linear registration of brains, a paradigm underlying most current neuroimaging analysis pipelines, Champollion V1 encodes individual morphological variability, which is essential for investigating genetic determinants of cortical folding.

To exploit the resulting high-dimensional, region-wise representations, we turn to multivariate GWAS methods, which were originally developed to aggregate association strength across correlated phenotype dimensions and are naturally suited to embeddings of this kind [23] (Fig. 1, step 3).

In the present study, we apply a multivariate GWAS framework based on Champollion V1 representations to 35,940 UK Biobank participants, with replication in 9,950 individuals from the ABCD cohort. We show that the resulting associations encompass and substantially enrich those recovered from conventional sulcal morphometric descriptors. Together, these analyses establish self-supervised representations of cortical folding as genetically informative phenotypes, extending conventional cortical morphometry while providing a general framework for evaluating representation learning through human genetics.

## 3 Results

### 3.1 Champollion V1 folding representations - associations

Self-supervised representations of cortical folding identify a substantially richer landscape of genetic associations than previously accessible through conventional morphometric phenotypes. Applying the Multivariate Omnibus Statistical Test (MOSTest) to the 56 regional self-supervised representations in 35,940 UK Biobank participants of white British ancestry, we identified 3,490 loci at the conventional genome-wide significance threshold (5 × 10*^−^*^8^) and 2,160 loci at a Bonferroni-corrected threshold accounting for the 56 regions tested (5 × 10*^−^*^8^*/*56 = 8×10*^−^*^10^), summed across regions (the same locus can be significant in more than one region). After merging redundant loci across regions (see Methods), these correspond to 567 and 328 independent loci, respectively. More than 90% of loci identified by the alternative MinP approach were also recovered by MOSTest (Extended Data). A detailed breakdown of associations by cortical region is provided in the Supplemental Material.

To assess whether learned representations capture genetic variation beyond conventional cortical morphometry, we performed multivariate GWAS using combinations of classical sulcal descriptors (length, surface area, mean depth, and maximum depth) for every sulcus within the same cortical regions. After merging loci across the 56 regions, this identified 162 independent loci at the conventional threshold (5 × 10*^−^*^8^) and 83 at the Bonferroni-corrected threshold (8 × 10*^−^*^10^), approximately threefold and fourfold fewer, respectively, than the 567 and 328 independent loci identified using self-supervised cortical folding representations. This nonetheless exceeds the loci reported by prior univariate GWAS of the same morphometric parameters (plus sulcal width) [24]. The Miami plot (Fig. 2a) shows that, at loci shared between the two approaches, association strength is consistently higher for Champollion V1 than for morphometric descriptors, while also revealing loci undetected by morphometry. Among the 567 independent loci identified from Champollion V1 representations, 188 independent loci have not been reported in the GWAS Catalog (March 2026). The Venn diagram (Fig. 2b) shows that 87% of loci identified using morphometric features are also captured by learned representations at the 5×10*^−^*^8^ threshold, with a similar pattern observed at 8 × 10*^−^*^10^. Furthermore, a scatter plot of log_10_(*p*) (Fig. 2c) indicates that the 22 loci detected by classical morphometry but not by the learned representations at the conventional threshold (highlighted in purple) all exhibit log_10_(*p*) values close to the significance threshold in the Champollion analysis. This is consistent with a thresholding effect rather than the absence of detectable genetic effects.

**Figure 2:**
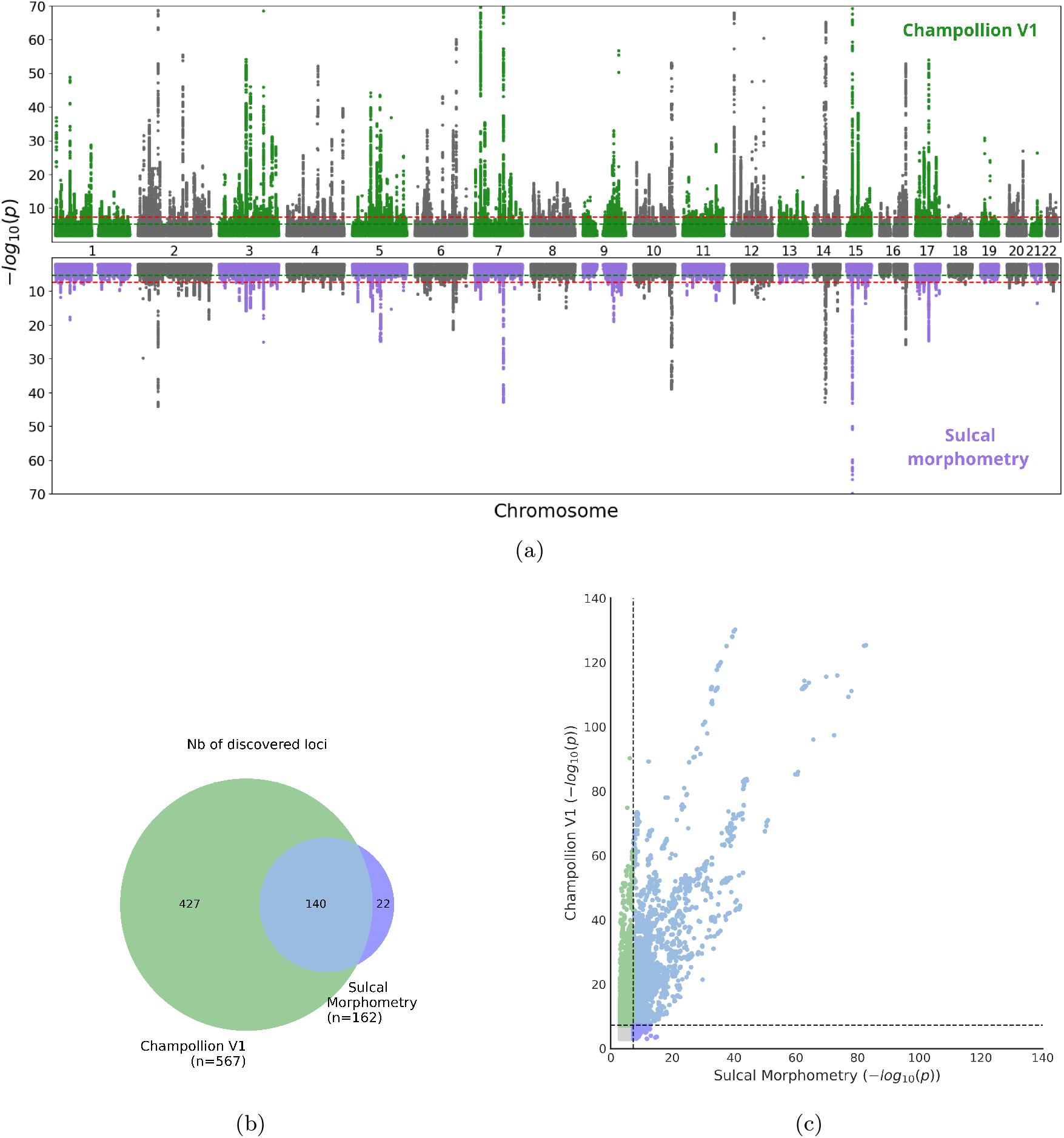
Champollion V1 representations subsume and extend the genetic associations identified by classical sulcal morphometric descriptors. **(a)** Miami plot comparing the strongest associations (using MOSTest) for all regions encoded by Champollion V1 on the 35,940 UKB white British ancestry subjects (green) to the strongest associations from the morphometric measurements (length of the sulci at the junction with the cortical hull, surface of the sulci, mean depth, and max depth) (purple). The conventional threshold of 5×10*^−^*^8^ is indicated with the red line. Note that the window is clipped for the log_10_(p-value) above 70. **(b)** Overlap of significant loci between Champollion V1 (green) and sulcal morphometry (purple). Overlap is considered to exist if there is at least one significant SNP within both loci. Numbers indicate the count of independent merged loci unique to each method or shared between them. **(c)** Scatter plot of SNP-based log_10_(*p*), with y-axis for Champollion V1 and x-axis for sulcal morphometry. Points above the diagonal indicate SNPs with stronger statistical evidence in Champollion V1 representations than in classical sulcal morphometry. The conventional threshold of 5 *×* 10*^−^*^8^ is indicated with the black dotted line. Note that *−* log_10_(*p*) are clipped at 140 on both axes.

Genetic associations were highly heterogeneous across the cortex. Among the learned-representation associations, the number of loci identified at the 5 × 10*^−^*^8^ threshold varied substantially across cortical regions, ranging from 130 for FPO-SCu-ScCal left to 11 for SFint-SR left (see Table 2 for nomenclature, and Supplemental Material for details). The number of loci per region was significantly correlated with the representation space SNP-based heritability estimated using LDSC (*R*^2^ = 0.74, *p* = 3 × 10*^−^*^17^; Fig. 3a). Regions with the highest heritability, i.e. primarily callosal, parieto-occipital, calcarine, posterior lateral, and sylvian (Fig. 3b), harboured the largest number of associations, whereas medial frontal regions, including SFint-SR (*h*^2^ *<* 0.10), showed the weakest associations. Region size did not explain either the number of loci or heritability. Indeed, regressing heritability against the number of voxels per region did not yield a significant association (*R*^2^ = 1.4 × 10*^−^*^5^, *p* = 0.98) (Fig. 3a). Notably, SFint-SR (57,171 voxels) is comparable in size to ScCal-SLi (48,048 voxels), yet exhibits approximately tenfold fewer significant loci at the 5 × 10*^−^*^8^ threshold.

**Figure 3:**
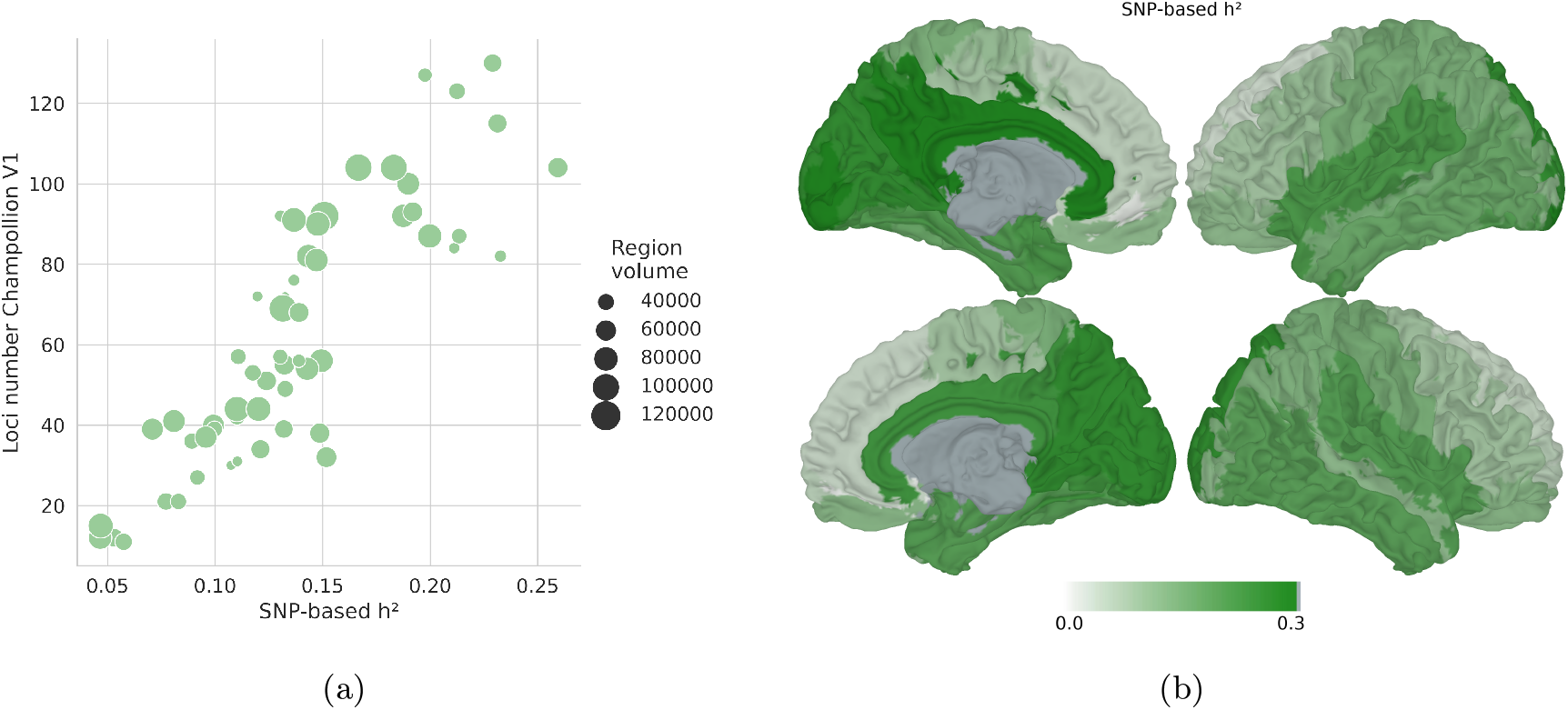
**(a)** Number of independent loci associated at *p <* 5 × 10*^−^*^8^ for each region encoded by Champollion V1 as a function of the SNP-based heritability of the representation (see Methods). Circle size represents the number of voxels in the bounding box of each region. **(b)** SNP-based heritabilities shown are the variance-weighted aggregate heritabilities across all 32 representation dimensions (see Methods). Colour scale: 0 (white) to 0.30 (dark green). Brain surface: MNI ICBM 152 white matter mesh (left lateral, left medial, right lateral, and right medial views).

Together, these results show that self-supervised representations not only recover and extend the genetic information captured by conventional descriptors, but also reveal that this information is concentrated in cortical regions of highest heritability.

### 3.2 Replication in the ABCD cohort

The 2,160 region-level loci identified in UKB discovery cohort (mean age 64.2 ± 7.7 years) at the Bonferroni-corrected threshold were tested for replication in the 9,950 participants of the ABCD cohort (mean age 9.9 ± 0.6 years, mixed ancestries; see Methods). These loci are retained at region-level resolution, rather than merged into independent loci, to enable the per-region replication analysis reported below. Merged across regions, they correspond to the 328 independent loci reported above. The UKB and ABCD cohorts differ substantially in age distribution, ancestral composition, and MRI acquisition protocol. This contrast constitutes a test of the extent to which the representations retain information about early neurodevelopmental processes. If the same genetic variants are associated with cortical folding morphology in both children (aged 8-11) and adults (aged 45-82), this would indicate that these representations capture morphological features established at least before 8 and preserved throughout life.

Of 2,093 lead SNPs identified at the 8 × 10*^−^*^10^ threshold in UKB and present in both cohorts, 1,079 replicated at the nominal threshold *α* = 0.05 in ABCD, corresponding to an overall replication rate of 51.6% (*SD* = 14.1% across regions) compared with a 5% rate expected under the null (Extended Data Table 4), which is higher than typical replication rates in imaging mGWAS (e.g. between 30-43% in Shadrin et al. MOSTest associations [25]). The best-replicated regions were STsbr right (71.4%), FIP-FIPPoCinf left (70.9%), and OCCIPITAL left (66.7%), all characterised by high heritability (see Table 2 for the sulcal nomenclature and Extended Data Table 3 for heritability). Conversely, the least-replicated regions, were SFmedian-SFpoltr-SFsup left (0/6 replicated lead SNPs) and SFint-SR left (0/4 replicated lead SNPs), precisely those with the lowest estimated heritability. A linear regression of the number of replicated loci on per-region heritability yields *R*^2^ = 0.71 (*p* = 3 *×* 10*^−^*^16^).

### 3.3 Gene-level enrichment analysis

Gene-level analysis revealed both broadly shared and region-specific genetic architectures. Among the genes identified across regions using MAGMA, a subset stood out for its ubiquity: *KANSL1*, *MAPT*, *STH*, *CRHR1*, *LRRC37A*, *ARL17B*, and *SPPL2C* were each associated with more than 40 of the 56 regions (at *p*_bonf_ *<* 0.05*/*19,264, to take into account the number of genes). These genes are all located within the 17q21.31 locus (chr17:43-45 Mb), a region of extended linkage disequilibrium that has been extensively associated with brain-related phenotypes [26, 27]. This locus was detected in 44 of the 56 regions tested, with *p*-values as low as 1.3 × 10*^−^*^54^, and was absent from the 12 least heritable frontal regions, consistent with the heritability gradient described above. It encompasses numerous genes previously implicated in cortical expansion (at least 29 genes), cortical curvature (4 genes), neurite density (13 genes), and cortical thickness (12 genes) as summarised in Warrier *et al.* [26].

Beyond this locus, several genes exhibited widespread association patterns across cortical regions, including *DAAM1* (chr14q23.1, 41 regions), *ZIC1* and *ZIC4* (chr3q24, 38 and 40 regions, respectively), *CCDC91* (chr12p11.22, 38 regions), *CENPW* (chr6q22.32, 37 regions), *C16orf95* (chr16q24.2, 36 regions), *SSH2* (chr17q11.2, 33 regions), *EPHA3* (chr3p11.1, 30 regions), and *CALCRL* (chr2q32.1, 27 regions). These genes have also been associated with a range of MRI-derived phenotypes, suggesting the presence of spatially distributed genetic contributions across the cortex. Detailed gene-level associations for each region are provided in the Supplemental Material.

Gene-level annotation using MAGMA, based on the 56 regional summary statistics, also showed a spatially patterned enrichment, as illustrated by *SEM1*, which is restricted to frontal regions, and *DACT1*, which is restricted to temporal regions (Extended Data Fig. 10a, 10b). The map obtained for *MAPT* (Fig. 4a) shows a spatial distribution that closely mirrors post-mortem regional expression profiles derived from the Allen Brain Institute [28], with enrichment in posterior parietal, lateral temporal, entorhinal, and precuneus regions (Extended Data Fig. 10d). This concordance between genetic associations and regional expression patterns is consistent with the hypothesis that the effects of *MAPT* on cortical folding morphology may be mediated by spatially patterned gene expression established during development and maintained into adulthood.

**Figure 4:**
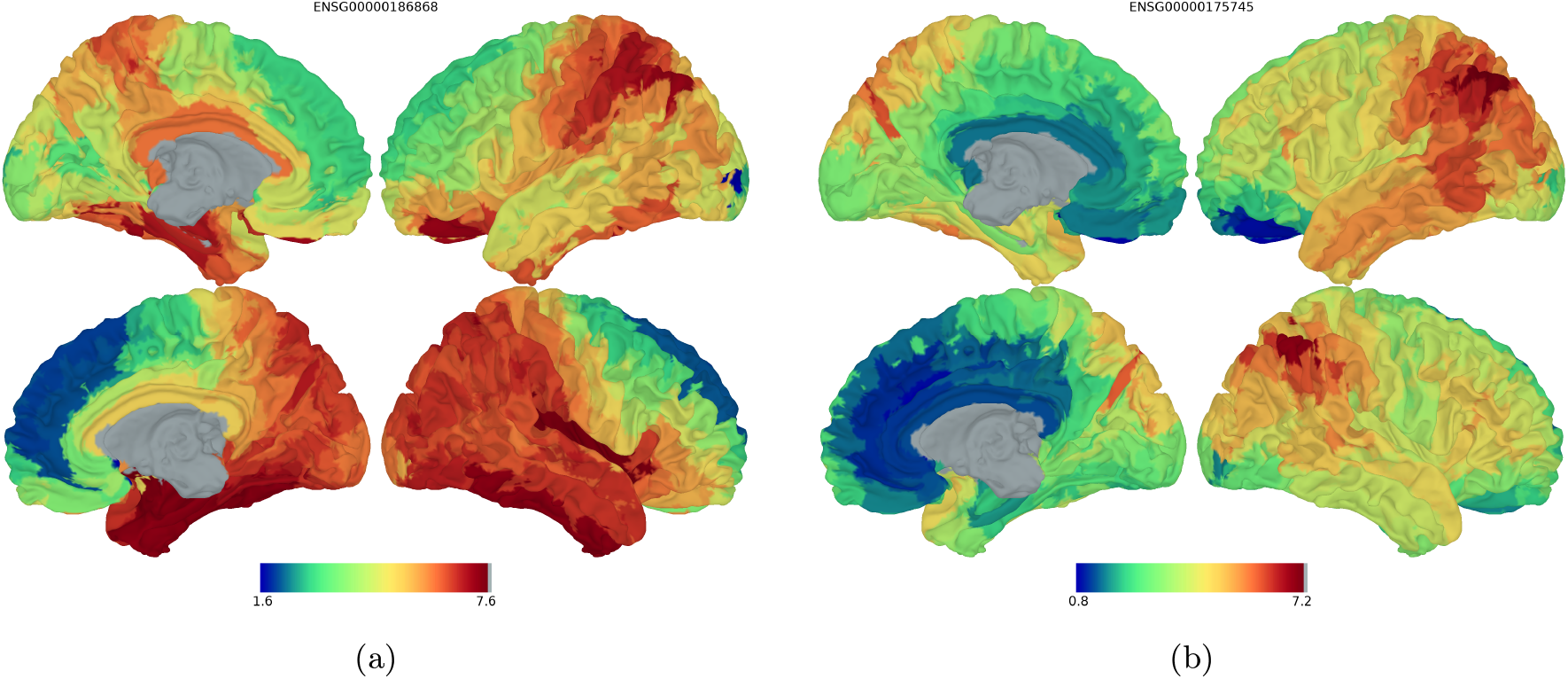
Cortex-wide MAGMA gene association maps for *MAPT* (panel **a**) and *NR2F1* (panel **b**). For each gene, MAGMA z-scores quantifying gene-level association with cortical folding morphology (encoded in Champollion V1 representations) are projected onto the cortical surface (MNI ICBM 152 white mesh, four views). Higher z-scores (warmer colours) indicate stronger association. *MAPT*-ENSG00000186868 (17q21.31) shows widespread posterior and temporal enrichment consistent with its Allen Brain Atlas expression profile [28]; *NR2F1* - ENSG00000175745 (5q14.3) shows a spatially restricted posterior enrichment consistent with its established role in controlling cortical gyrification [31].

In contrast, the association map for *NR2F1* (Fig. 4b) exhibits a spatially restricted pattern, primarily localized to the posterior parietal cortex, in strong agreement with the known biology of this transcriptional regulator. *NR2F1* (also known as COUP-TFI) follows a high postero-lateral to low rostro-medial expression gradient in cortical progenitors and post-mitotic neurons [29], a pattern conserved in humans [30]. While the effect sizes of common *NR2F1* variants in the general population are far smaller than those of haploinsufficiency in the context of Bosch-Boonstra-Schaaf optic atrophy syndrome (BBSOAS), the spatial concordance of our associations with the cortical territories affected in syndrome patients provides convergent evidence that *NR2F1* influences posterior cortical morphology in a dose-dependent manner [31, 32]. The spatial association pattern identified here therefore provides population-genetic support for a role of *NR2F1* in regional variation of posterior cortical folding.

Together, these findings indicate that learned representations not only increase statistical power for locus discovery but also reveal spatially organised biological effects that are consistent with early neurodevelopmental programs.

### 3.4 Gene-set enrichment analysis

Because the 56 sulcal regions partially overlap anatomically and are therefore not statistically independent, gene-set-level associations were combined across regions using a Stouffer statistic corrected for their empirical inter-region correlation structure, yielding a single global z-score per gene set (see Methods). Neurodevelopmental gene sets were identified as the strongest enrichments across cortical folding representations (Table 1). The most significantly enriched gene sets (regulation of anatomical structure morphogenesis, generation of neurons, and neurogenesis) (*z*_Stouffer_ *>* 6.9, *p_bonf_ <* 5 × 10*^−^*^9^) implicate biological processes involved in neuronal production, tissue morphogenesis and cortical development during fetal life. For example, neurogenesis peaks during the second trimester [2], coinciding with the emergence of primary sulci. These enrichments align with prior GWAS of cortical thickness and surface area [23, 33], suggesting that Champollion V1 captures underlying genetic effects acting during a shared prenatal window when both regional expansion and folding patterns are established. Unexpectedly, gene sets related to mesenchymal cell differentiation and mesenchyme development (*z*_Stouffer_ = 7.0 and 6.9, respectively) emerged as strongly associated with cortical folding. This point is addressed in the discussion.

**Table 1:**
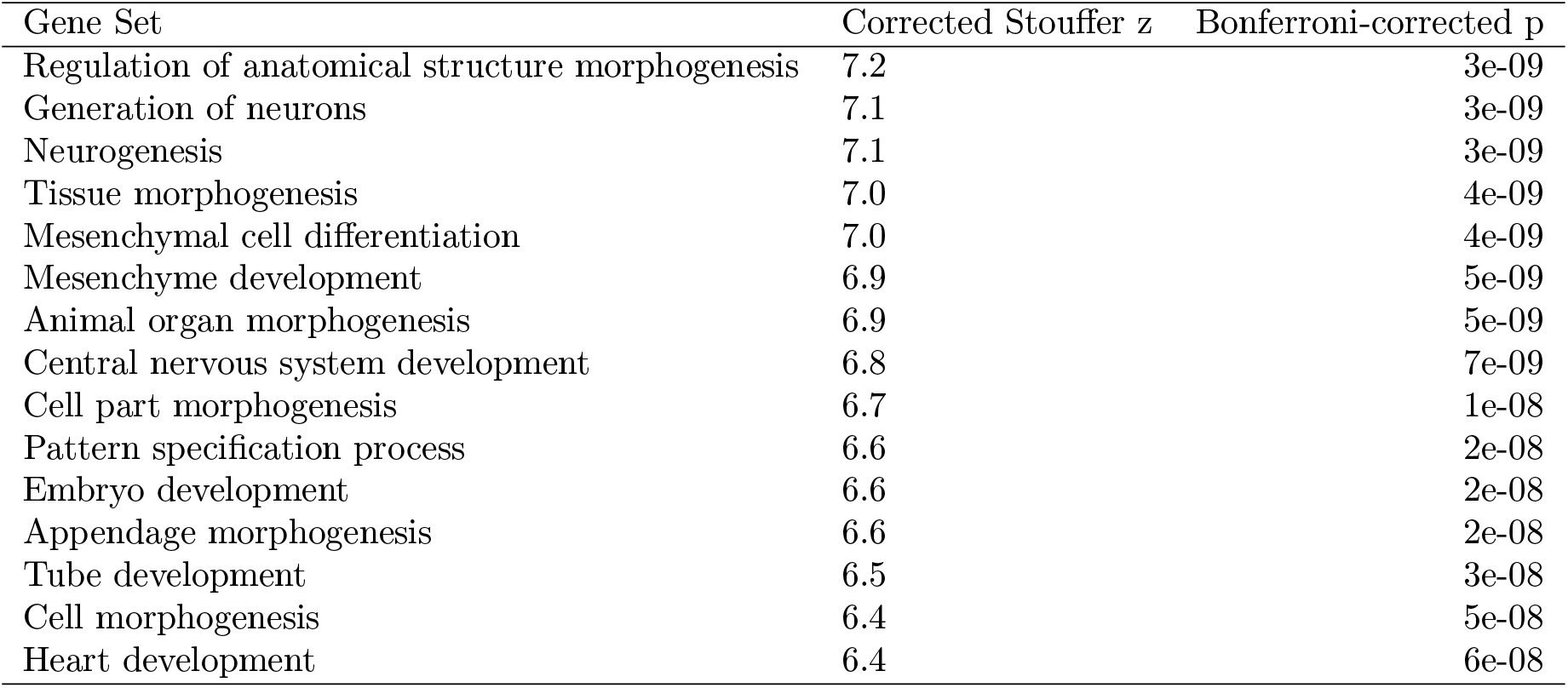
Gene-set analysis with MAGMA. Top 15 gene sets associated across the 56 sulcal regions, after a corrected Stouffer analysis.

**Table 2:**
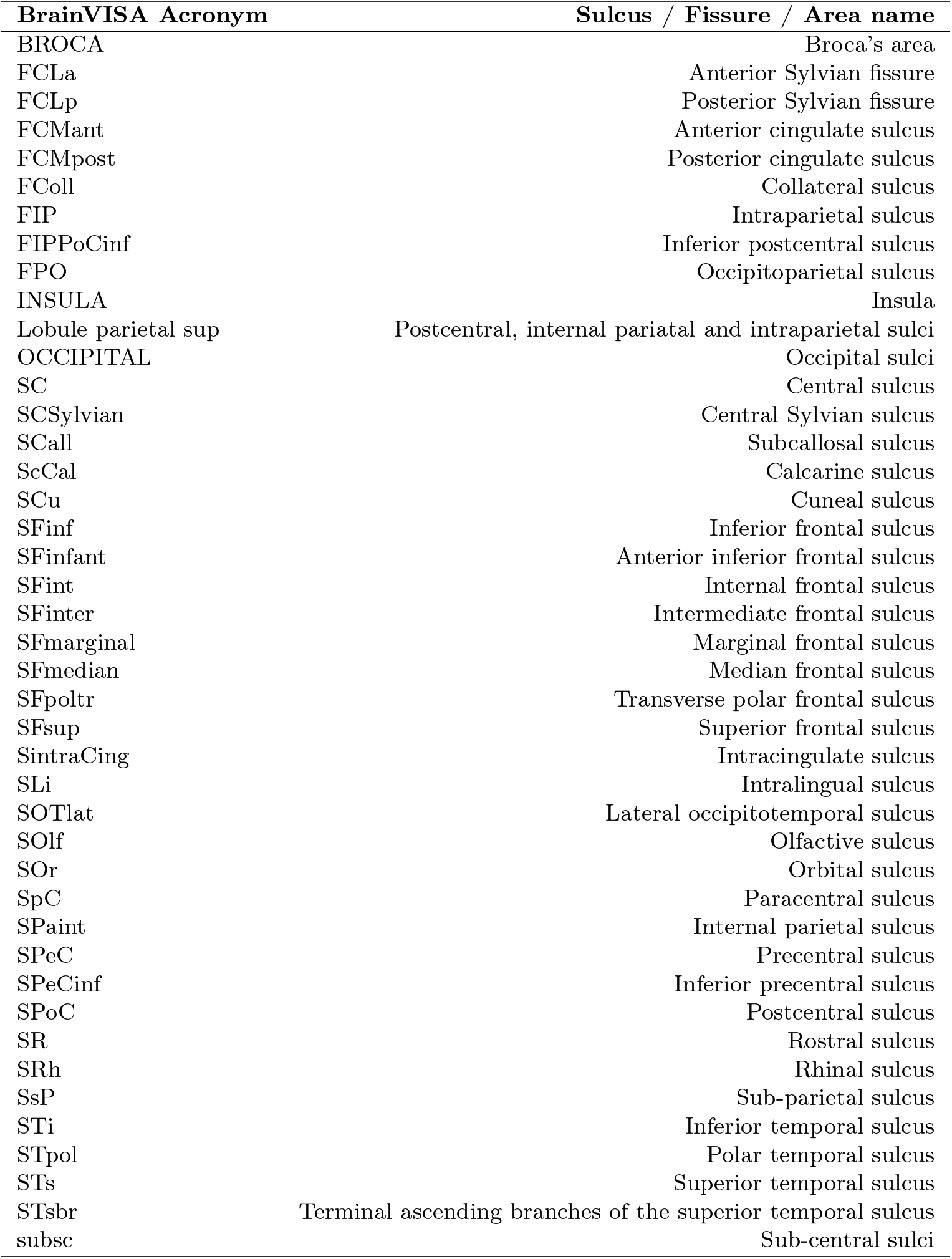
Acronyms for sulcus, fissure, and areas names.

### 3.5 BrainSpan enrichment analysis

Gene expression enrichment analysis using BrainSpan revealed a temporally specific developmental profile. Combined across the 56 regions using the Stouffer framework, genes associated with cortical folding morphology showed strong enrichment for expression during fetal stages, spanning 8 to 37 post-conception weeks (*z*_Stouffer_ between 3.5 and 9.0, Bonferroni-corrected *p <* 6 × 10*^−^*^3^), with peaks at 9 and 16 post-conception weeks. These stages correspond to periods of active progenitor proliferation, neurogenesis, and neuronal migration [2].

This enrichment rapidly diminishes after birth, becoming undetectable by 4 months postnatally, and re-emerging only marginally at 2 years (*z* = 1.9) (Fig. 5). This predominantly prenatal expression profile is consistent with the long-term stability of cortical folding patterns across adult life [11, 12].

**Figure 5:**
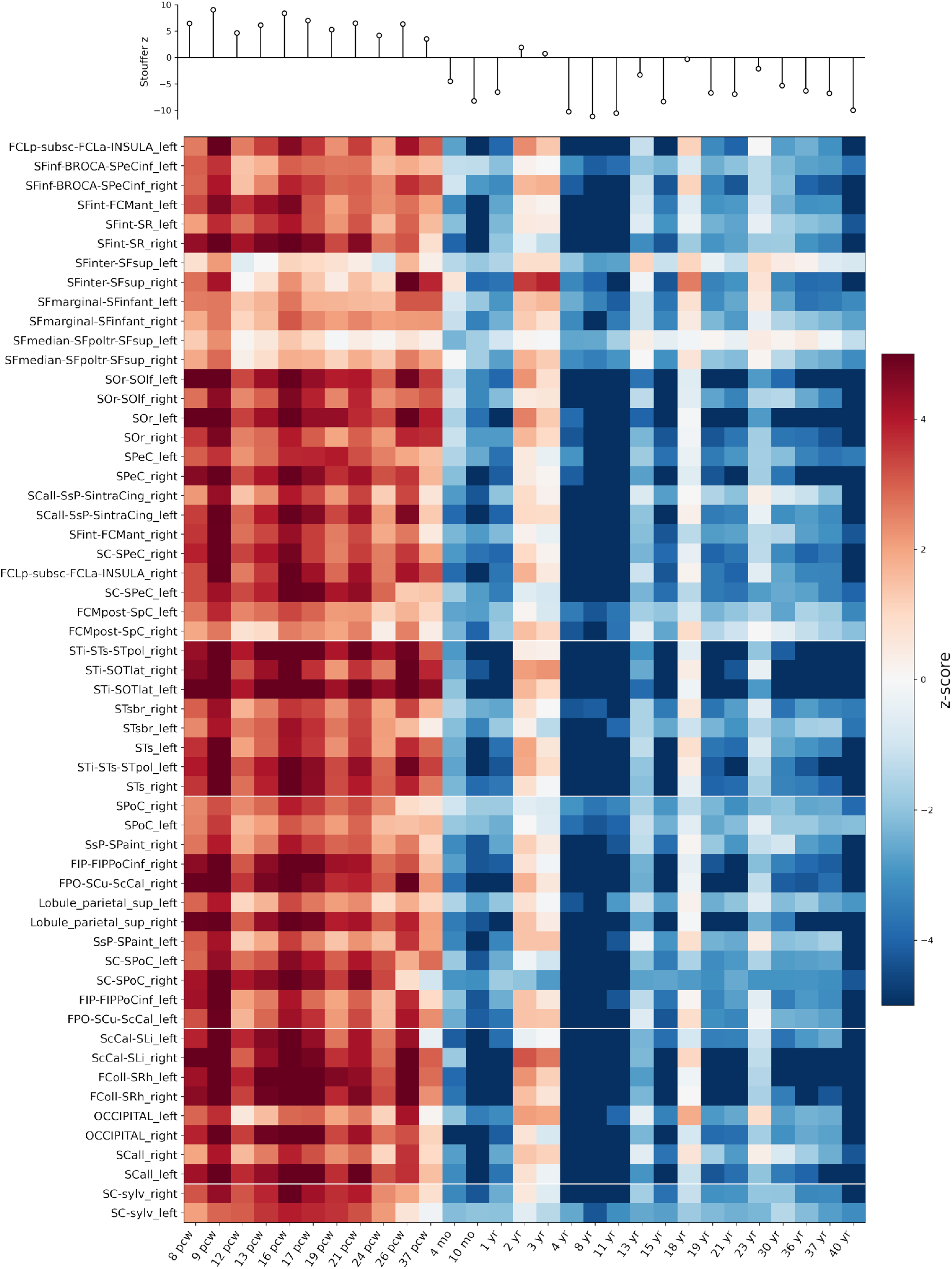
Heatmap of MAGMA results showing enrichment of genes across 56 sulcal regions (rows) and 21 BrainSpan developmental stages (columns). Colours represent z-transformed p-values (red: higher-than-expected expression; blue: lower). Stages are ordered chronologically (pcw: post-conception week; mo: month; yr: year). Regions are grouped by lobe, separated by the thin white lines (Frontal, Temporal, Parietal, Occipital/Calcarine, Lateral fissure, Central sulcus). The upper panel shows the Stouffer meta-analysis z-score combining the corrected evidence across all 56 sulcal regions at each developmental stage (vertical lollipop plot), summarizing the overall strength of enrichment. Enrichment is positive when the Stouffer z is positive. The enrichment peaks at 9 and 16 pcw.

### 3.6 Single cell enrichment analysis

By mapping the cortical regions used in the Champollion V1 framework to the major cortical regions defined in Bhaduri *et al.* 2021 [34], we investigated whether the sets of genes associated with each self-supervised learned representation were preferentially enriched in specific fetal brain cell types across developmental time. We selected two highly heritable regions, left SCall-SsP-SintraCing (*h*^2^ = 0.26 ± 0.01) and right FPO-SCu-ScCal (*h*^2^ = 0.23 ± 0.01), to illustrate the dynamics observed across regions (Figure 6).

**Figure 6:**
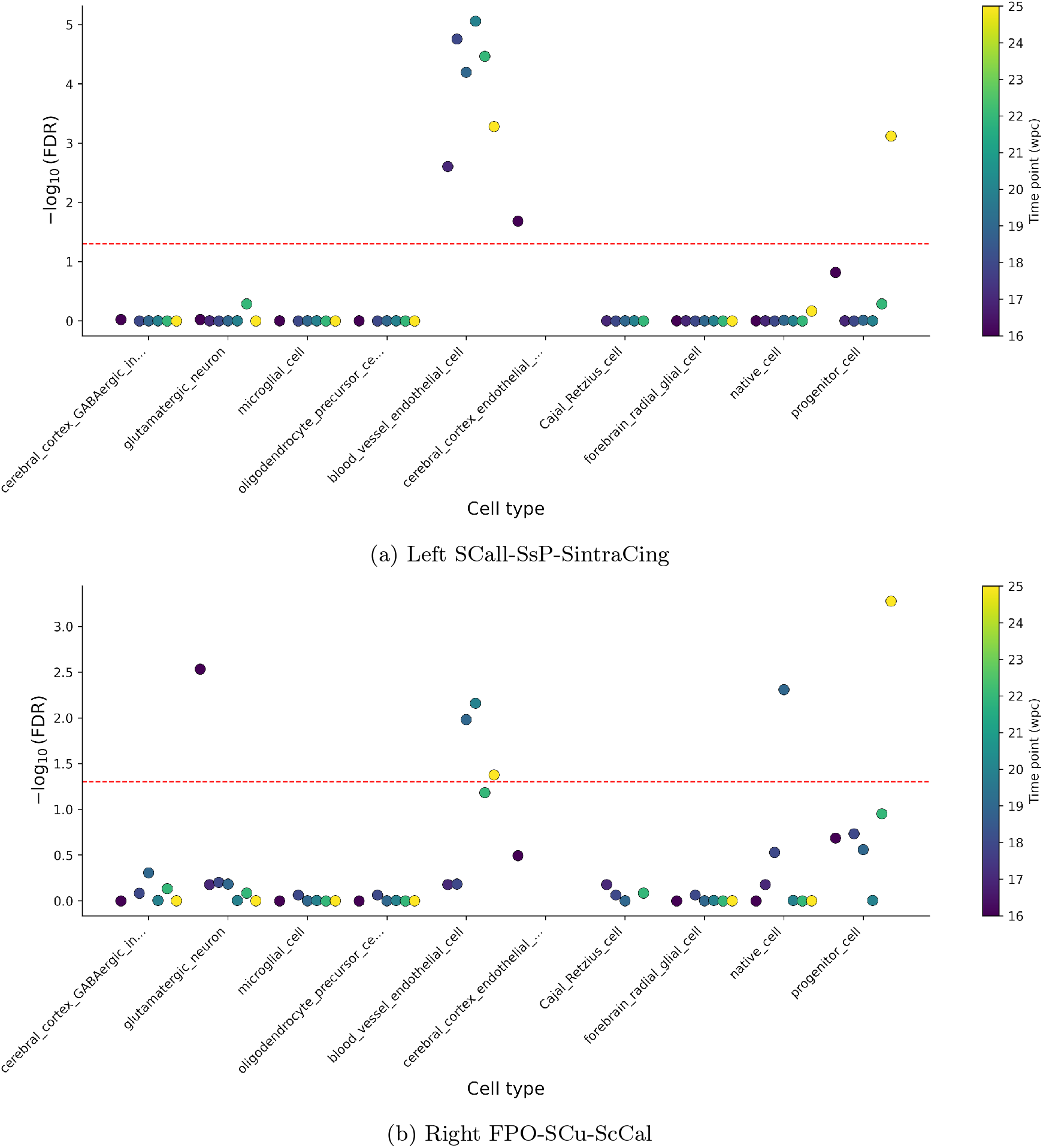
Cell-type enrichment across developmental time points for the **(a)** left SCall-SsP-SintraCing and the **(b)** right FPO-SCu-ScCal regions. Each point represents the enrichment significance of a given cell type obtained from the MAGMA gene-property analysis using fetal cortical single-cell RNA-sequencing data. The y-axis shows the FDR-corrected significance (-log_10_(FDR)), where larger values indicate stronger evidence for enrichment after Benjamini-Hochberg correction across the cell types tested within each developmental time point. The -log_10_(0.05) threshold is indicated with the red line. Point colours denote the developmental stage (post-conception weeks, PCW), allowing visualisation of temporal changes in the cell-type enrichment profile throughout cortical development.

Across the 16-22 pcw window, the strongest enrichment was observed for genes specifically expressed in blood vessel endothelial cells (*log*_10_(FDR) = 5). Endothelial cells play an essential role in the establishment of the cerebral vasculature during corticogenesis, providing the vascular network required to sustain the developing cortex. At the last available fetal stage (25 pcw), enrichment additionally emerged for progenitor-cell gene expression (*log*_10_(FDR) = 3), suggesting a shift toward genes expressed in neural progenitor populations as development progresses.

Together, these analyses suggest that the genetic architecture captured by learned cortical folding phenotypes maps onto distinct cellular programs across fetal corticogenesis.

### 3.7 Genetic correlations with psychiatric disorders

Genetic correlations between the Champollion V1 representation spaces and seven psychiatric and neurological disorders (schizophrenia, bipolar disorder, attention-deficit/hyperactivity disorder (ADHD), autism spectrum disorder (ASD), depression, neuroticism, and post-traumatic stress disorder (PTSD)) were estimated using LDSC correlations (see Methods) corrected for the number of tests 56 *×* 7 (*p*_bonf_ *<* 1.28 *×* 10*^−^*^4^).

For schizophrenia, three regions survived the full Bonferroni correction: right SC-SPeC (*p* = 9 × 10*^−^*^7^), right SFint-FCMant, also known as the cingulate cortex (*p* = 4 × 10*^−^*^5^), and right SC-sylv (*p* = 1 × 10*^−^*^4^). Two additional regions also encompassing the right central sulcus showed convergent correlations failed to survive correction: right SC-SPoC (*p* = 6 × 10*^−^*^4^), and right SPeC (*p* = 7 × 10*^−^*^4^), suggesting that the correlation is carried by the morphology of the right central sulcus as a whole (Extended Data Fig. 11). Strikingly, no region in the left hemisphere showed even nominal association with schizophrenia, despite the strong inter-hemispheric symmetry of the genetic associations described in the Genetic Dice similarity across regions section in Extended Data. This hemispheric specificity of the schizophrenia association suggests that lateralized cortical folding morphology, rather than globally symmetric folding architecture, may be a mediating neurobiological substrate of genetic risk for schizophrenia. This is consistent with reports of cortical asymmetry in schizophrenia [35] and with the known genetic overlap between handedness and psychosis risk [36].

A closer examination of the representation space encoding the right SC-sylv region revealed that the correlation was primarily driven by the second principal component (*r_g_*= − 0.14 ± 0.03, *p* = 3× 10*^−^*^5^). Visualization of this component using the decoder [37] showed that it encodes the size of a knob-like structure within the right central sulcus (Extended Data Figure 12). Interestingly, the size of this knob in the right hemisphere has previously been associated with left-handedness, with left-handed individuals exhibiting a larger knob on average than right-handed individuals [38]. Causal inference would require Mendelian randomisation analyses, which are beyond the scope of this study.

For bipolar disorder, no region survived the full Bonferroni correction, although right SC-SPoC showed the strongest correlation (*p* = 1 × 10*^−^*^3^), comparable to ScCal-SLi left (*p* = 1 × 10*^−^*^3^) and right SC-sylv (*p* = 7 × 10*^−^*^3^). The latter was also the second region most strongly associated with schizophrenia, consistent with the high genetic correlation reported between these two disorders (*r_g_* = 0.70 [39]). The interpretation of the association between bipolar disorder and the ScCal-SLi left region (calcarine-intralingual sulcus) remains unclear and warrants further investigation.

For PTSD, the right SFmedian-SFpoltr-SFsup region showed a significant correlation surviving full Bonferroni correction (*p* = 5 × 10*^−^*^6^). Neuroticism and ADHD showed nominal, sub-threshold associations with the right FCMpost-SpC (*p* = 2 × 10*^−^*^4^) and right SC-SPeC (*p* = 6 × 10*^−^*^4^) regions, respectively, that did not survive Bonferroni correction. No region showed even nominal association with the remaining disorders (autism spectrum disorder, depression).

## 4 Discussion

A central challenge in imaging genetics is that conventional phenotypes necessarily compress complex anatomical structures into a small number of handcrafted measurements. By combining self-supervised representations of cortical folding with multivariate GWAS, we show that higher-dimensional phenotypes substantially increase genetic discovery while remaining biologically interpretable. Learned representations recovered nearly all loci identified using classical sulcal morphometry and revealed hundreds of additional associations, demonstrating that they capture genetically informative variation beyond conventional descriptors.

A key result of the present work is that the associations obtained with Champollion V1 largely encompass those identified using the same multivariate framework applied to conventional sulcal morphometric descriptors (length, surface area, mean depth and max depth), within the same cortical regions and cohort. More than 87% of morphometric loci were recovered among the Champollion V1 loci, while the remaining 13% of loci appeared to reflect threshold effects rather than a complete absence of detectable genetic effects. More broadly, this supports the idea that the cerebral cortex can be investigated using higher-dimensional phenotypes that are richer, if less immediately interpretable, than conventional measures, without sacrificing the associations already established through those measures.

This extension toward higher-dimensional phenotypes nevertheless warrants an important caveat. Detecting a larger number of loci does not necessarily imply that more biologically meaningful information is being captured, particularly when the exact nature of the encoded phenotype is not fully controlled. In the present study, this limitation is mitigated by the extensive preprocessing applied to the MRI data, including cortical segmentation and extraction of cortical folding skeletons, such that the representation spaces of Champollion V1 specifically encode cortical folding patterns. Unlike approaches that learn representations directly from minimally processed MRI scans, which often lack clear anatomical interpretability, the present framework retains a well-defined morphological meaning related to the shape and spatial organisation of cortical folds. Nevertheless, the proportion of variance explained by genetics, as estimated through SNP-based heritability ranging from 0.05 to 0.26, represents only a modest fraction of the total variance captured within these representation spaces. The representation dimensions also encode non-genetic sources of variability, including environmental and stochastic effects, whose characterisation constitutes an important direction for future work.

Regional differences in SNP-based heritability further indicate that the genetic architecture of cortical folding is heterogeneous across the cortex. Frontal regions consistently exhibited the lowest heritability, consistent with prior reports [40], whereas callosal, calcarine and posterior regions showed the strongest heritability. This pattern is consistent with previous observations that earlier-forming folds are generally under stronger genetic control than later-developing cortical regions [7].

Replication across the UK Biobank and ABCD cohorts, despite substantial differences in age, ancestry, and MRI acquisition, indicates that the traits captured by learned cortical folding representations are stable across the lifespan. This developmental interpretation is further supported by the enrichment of genes involved in neurogenesis, neuronal differentiation, and tissue morphogenesis, together with their preferential fetal expression in BrainSpan. Moreover, associations obtained from the learned representations are largely orthogonal to those obtained from sulcal opening (see Extended Data), a measure that is itself lifespan-dependent. Together, these results support the hypothesis that secondary and tertiary sulci are biomarkers of early neurodevelopment, and that the representations learned by Champollion V1 successfully capture their variability. This makes these representations particularly well suited to the study of neurodevelopmental disorders.

Two gene sets nevertheless stood out across multiple cortical regions because of their unexpectedly strong association with cortical folding morphology: mesenchymal cell differentiation and mesenchymal cell development (*z*_Stouffer_ = 7.0 and 6.9, respectively). Several biological mechanisms may account for this finding. Interactions between neural crest-derived mesenchymal-like cells and cortical development have previously been documented, notably through growth factor signaling pathways [41]. In addition, mesenchymal cells may contribute to the formation of the pericortical extracellular matrix (ECM), which has been proposed to regulate cortical progenitor signaling and cortical folding mechanics [10, 42]. The enrichment of these pathways in our results may therefore reflect a contribution of the pericortical mesenchymal environment to the mechanical constraints shaping cortical gyrification, a developmental dimension that remains largely unexplored in human genetics. More broadly, variation in mesenchymal differentiation could simultaneously influence craniofacial morphology [43] and the biomechanical constraints applied to the developing cortex, thereby indirectly affecting folding patterns. Consistent with this interpretation, the *WP_NEURAL_CREST_DIFFERENTIATION* gene set reached significance in 5 out of the 56 cortical regions analysed (FColl-SRh left, SC-sylv right, STi-STs-STpol left, STs left, SsP-SPaint right). However, further experimental validation is needed to confirm this hypothesis.

Several limitations should nevertheless be acknowledged. First, the discovery analyses were restricted to participants of white British ancestry in the UK Biobank, and the usage of mixed ancestries in the ABCD cohort may reduce replication rates, but was still preferred for statistical power over ancestry-specific replication analysis. Indeed, the 5,269 participants of European ancestry in ABCD did not provide sufficient power for replication. Second, the present study focused exclusively on autosomal variation. Sex chromosomes were excluded because of their distinct inheritance structure and analytical complexity. In addition, the present analyses focused on common genetic variation captured through standard GWAS approaches. Rare variants, structural variants, and other forms of genetic variation remain largely unexplored in the context of cortical folding morphology and may contribute substantially to inter-individual variability. Because the Bhaduri atlas spans fetal development only up to 25 pcw, it is not possible to determine from these data whether this developmental trajectory persists into the third trimester or postnatal development.

Beyond cortical folding specifically, these findings suggest that representation learning offers a general strategy for constructing genetically informative phenotypes from complex biomedical images.

## 5 Methods

### 5.1 Datasets

Two independent datasets were used in this study. First, the UK Biobank dataset (application number 64984), with imaging and genotyping data from 42,433 participants. The final sample for the study included 35,940 participants of white British ancestry (47.5% male), aged 45 to 82 years (UK Biobank field: 53, first imaging visit), with a mean age of 64.2 ± 7.7 years.

Second, data were drawn from the Adolescent Brain Cognitive Development (ABCD) study, with imaging and genotyping data from 11,159 participants. The final sample included 9,950 participants from release r6 (52.4% male, mean age 9.9 ± 0.6 years, range 8.3 to 11.3 years) including 5,269 European ancestry, 1,460 African ancestry, 2,021 Hispanic, 195 Asian ancestry, and 1,005 other ancestries. Only data from session 00A were included to avoid multiple time points per subject.

### 5.2 Imaging data processing

The Morphologist v5.0 pipeline from the BrainVISA software suite [44] was applied to 42,433 T1-weighted images from the UK Biobank (field: 20252, first imaging visit) and the Morphologist v6.0 to the 11,159 T1-weighted images from the ABCD cohort (session 00A), both acquired at 1 mm isotropic resolution. Note that Morphologist v5.0 and v6.0 use the same segmentation pipeline, with no differences in the code. Those are the initial imaging datasets before Quality Control (QC) and multimodal matching. For each hemisphere, this processing pipeline generates a 1 mm-resolution skeleton of the cortical folds in native space, representing a negative cast of the cortex [15, 16]. This skeleton captures cortical folding patterns as a binary volumetric image with a maximum thickness of one voxel (step 1, Fig. 1).

Because this representation does not encode variations in cortical thickness or sulcal opening, it enables the analysis to focus exclusively on folding patterns, which are generally considered to be more stable features of cortical morphology across lifetime [11, 12]. The skeleton was subsequently affinely normalised to the MNI ICBM 2009 space and then downsampled to a 2 mm isotropic resolution, which is sufficient to capture the principal folding patterns [8].

### 5.3 Quality control

Participants with poor Morphologist segmentation (BrainVISA software), such as missing structures, missing graph or invalid file format, were excluded based on automated QC procedures. Quality control was performed using automated criteria; individual-level visual inspection was not applied. This procedure was applied consistently to the UK Biobank cohort with the Morphologist v5.0 and to the ABCD cohort with the Morphologist v6.0.

To further refine the quality of cortical folding skeletons, 18 participants with an abnormally high number of voxels in the skeleton were excluded. Specifically, individuals whose total voxel count exceeded 1.2 times the 9th decile of the voxel count distribution within each cohort were removed. Visual inspection confirmed that all such cases exhibited clear defects. This threshold was determined empirically.

QC was performed blind to downstream analyses.

### 5.4 Definition of cortical regions

Regions of interest were defined using the BrainVISA sulcal atlas (Extended Data Fig. 7), with the aim of capturing anatomically meaningful units corresponding to major sulci or gyri. This atlas is derived from the PCLEAN database [15], which is based on manual sulcal annotations.

For each sulcus, the smallest bounding volume enclosing all its occurrences across subjects from the PCLEAN database was estimated in MNI space. Groups of sulci were then manually organised around major anatomical landmarks, including principal sulci, and major gyri. For each such group, the corresponding regional volume was defined as the aggregation of the bounding volumes of the sulci composing the region. These volumes were subsequently dilated by 5 mm in all directions to produce masks that robustly encompass the associated sulci and gyri, especially for subjects that are different from the PCLEAN database. From this stage onward, explicit sulcal labeling of individual brains is no longer required to apply the regional masks. This procedure defines the Champollion V1 atlas used throughout the study, and can be applied with the Cortical Tiles tool (https://github.com/neurospin/cortical_tiles.git).

For each subject, the cortical folding skeleton of each hemisphere was masked according to the regions defined by the Champollion V1 atlas [8]. Each hemisphere was subdivided into 28 partially overlapping regions, yielding a total of 56 regions across the whole brain. Region sizes vary substantially (for instance, from 26 ± 34 ± 23 voxels for S.Or. right to 34 ± 67 ± 57 voxels for S.T.i.-S.T.s.-S.T.pol.), as does the shape complexity of the folding patterns they contain.

Because regions are allowed to overlap, folding patterns may contribute to multiple regions.

### 5.5 Multivariate phenotype 1: Champollion V1 folding representations

Pre-trained contrastive models, independently fitted for each cortical region, separately per hemisphere, provide a 32-dimensional representation space per region [8]. The training took place on 42,000 subjects from the UK Biobank dataset. Note that no genetic labels were used for training nor optimisation. These representation spaces are designed to compactly encode the shapes of cortical folding patterns, capturing information related to fold length, depth, surface area, curvature, and relative spatial organisation.

For a given cortical region, folding morphology is thus represented as a 32-dimensional parametric embedding that captures the principal modes of shape variation observed across individuals (Step 2.a Fig. 1). As previously demonstrated [8], these representation dimensions show significant associations with conventional morphometric descriptors, as assessed through regression and classification tasks on folding patterns in orbito-frontal, central, and intra-parietal regions.

To facilitate downstream analyses, principal component analysis (PCA), fitted on the UK BioBank dataset, was applied to each regional compressed representation capturing sulcal shapes, to obtain orthogonal components ordered by explained variance. All components are retained for analysis. PCA loadings estimated in UK Biobank were applied to the ABCD cohort without refitting.

### 5.6 Multivariate phenotype 2: BrainVISA morphometric parameters

Following the BrainVISA nomenclature [44], an automated deep learning-based sulcal labeling model from Morphologist v5.0 [15] was used to segment the cortical folding skeletons into individual sulci. To ensure sufficient sample size and robustness, only sulci present in more than 90% of individuals were retained, yielding a sample size comparable to that of the Champollion V1 representations (~35,000 subjects).

For each retained sulcus, the following morphometric parameters were extracted: geodesic length at the junction with the cortical hull, surface area, mean depth, and maximum depth. These measurements were computed in Talairach space rather than native space to ensure consistency with the Champollion inputs. The MNI ICBM 2009 space used in Champollion and the Talairach space used in BrainVISA can be considered equivalent up to an affine transformation.

Sulcal opening and cortical thickness were not included in the multivariate phenotype, as these measures are not directly defined in cortical folding skeletons, which represent medial surfaces without thickness. However, sulcal opening on its own was studied apart to show orthogonality of the associations compared to those from the representation space (see Extended Data Fig. 8).

To ensure consistency with the Champollion V1 regional framework, sulci morphometric parameters were concatenated according to the same regional definitions. Indeed, a Champollion V1 region was defined as the aggregation of the bounding volumes of the sulci composing the region. Thus, the morphometric characterisation of a given region is based on the set of length, surface, and depth measures associated with all sulci that were used at the definition step to compose that region (Step 2.b Fig. 1). Because regions are partially overlapping, a given sulcus may contribute to multiple regions.

### 5.7 Genetic data processing

Genotyping data were processed in parallel with imaging data. For the UK Biobank cohort, among the 42,433 individuals with imaging data, 41,395 had valid genotyping data (GRCh37), including 35,940 participants of white British ancestry (UK BioBank field 22006) retained for the main analyses.

For the ABCD cohort, to maximise statistical power, genetic analyses were conducted on all individuals with available genotyping data (GRCh38), yielding a total sample size of 9,950 subjects. The ABCD cohort was used exclusively as an independent replication sample. No ancestry-specific replication analysis was conducted, since 5,269 subjects would not give enough statistical power to replicate all the significant SNPs.

The present association analyses were restricted to autosomal variants; sex chromosomes (X and Y) were excluded.

Imputed genotype data were used for both cohorts: UK Biobank data corresponded to imputation release version 3 [18], while ABCD data were imputed using the TOPMed reference panel. The same quality control filters were applied to both cohorts, resulting in 8,109,955 SNPs for UK Biobank and 8,961,447 SNPs for ABCD.

Genotype quality control was performed using PLINK2 (version 17-02-22). Only autosomal, biallelic single nucleotide polymorphisms were retained (–snps-only, –max-alleles 2). Variants with a minor allele frequency (MAF) below 1% were excluded (–maf 0.01). Standard quality control filters were applied at both the variant and individual levels (–geno 0.1, –mind 0.1), and deviations from Hardy-Weinberg equilibrium were filtered using a significance threshold of *p <* 1 *×* 10*^−^*^9^ (–hwe 1e-9).

### 5.8 Multivariate genetic associations

Pre-residualization was performed separately for each cohort, using a linear regression model implemented in statsmodels.

(a) For the UK Biobank cohort, the following covariates were included: sex (field 31), age at imaging (field 53), age^2^, age × sex and age^2^ sex interactions, imaging center (field 54), total intracranial volume (TIV) using CAT12 (v12.8), and the first 20 genetic principal components (field 22009). For each dimension of the 32-dimensional representation space, across all 56 regions, we regressed subject-level coefficients (35,940 white British ancestry participants) against each of the 40 leading genetic principal components, yielding 32 × 56 p-values per genetic PC. This tests whether the self-supervised representations nonetheless carry structure aligned with population genetic axes (Supplemental Material Fig.1). Seven genetic PCs (PC4, 5, 9, 10, 11, 12, 14) show a modest excess of low p-values relative to the uniform expectation, and were regressed out at the pre-residualisation step, as the first 20 genetic PCs were included in the covariates. For the ABCD cohort, equivalent covariates were used: age at first visit (ab_g_dyn_visit_age), age^2^, sex (ab_g_stc_cohort_sex), the first 20 genetic principal components (gn_y_popstruct_pc_), scanner serial number (ab_g_dyn_design_mr_serial), and TIV estimated using CAT12 (v12.8).

All measures were residualized with respect to total intracranial volume to specifically isolate regional morphological effects. This procedure removes global contributions related to overall brain size, which may otherwise influence folding shapes through the relationship between brain volume and gyrification.

(b) Each regional representation space was treated as a multivariate phenotype. SNP-based heritability 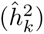 was estimated independently for each dimension using Linkage Disequilibrium Score Regression (LDSC). The number of dimensions with 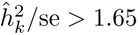 (approximately corresponding to a one-sided *p <* 0.05) varied across cortical regions, ranging from 16 (SFint-SR left) to 31 (STi-SOTlat left), with a mean of 26.5 ± 3.4. To avoid bias from selecting only significantly heritable dimensions, all 32 dimensions were retained for downstream genetic analyses.

To summarise regional heritability as a single scalar comparable across regions, an aggregated multivariate heritability measure was defined. Because the 32 representation dimensions are orthogonal by construction after PCA, the aggregated heritability can be computed as a variance-weighted sum of per-dimension heritabilities: 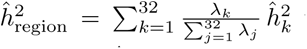 where *λ_k_* denotes the variance explained by the *k*-th principal component and 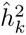 its LDSC-based heritability estimate. The weights 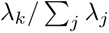 correspond to the proportion of variance explained by each component. Because all components are retained, the weights sum to one and no truncation bias is introduced. The associated standard error was derived by propagation under the assumption of independence across dimensions: 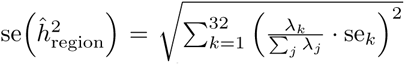. The advantage of this heritability is to take into account the proportion of the explained variance for each component, contrary to the heritability defined in [45], which would average non comparable components.

(c) Multivariate genetic associations were assessed using the Multivariate Omnibus Statistical Test (MOSTest), which combines summary statistics from univariate genome-wide association studies (GWAS) conducted on each phenotype. MOSTest was selected over MinP as it has been shown to provide greater statistical power [23]. For each SNP, univariate GWAS z-scores across the pre-residualized dimensions of a given regional representation were combined into a multivariate test statistic defined as *χ*^2^ = *z_j_R*^−1^ *z_j_^T^*, where *z_j_*is the p-dimensional row vector of z-scores (where p = 32, the number of principal components) and *R* is the *p × p* empirical correlation matrix of the test statistics under permutation of the genotypes. To preserve the correlation structure across phenotype dimensions, genotype permutations were performed once per SNP, and the resulting permuted genotype vector was used to compute association statistics across all phenotype dimensions. Because a PCA was made, *R* = *Id* (no correlation between the phenotype dimensions). The null distribution of the test statistic was estimated from these permutation-derived statistics. Rather than relying on the theoretical chi-square distribution, a Gamma distribution was fitted to the empirical distribution of the test statistic under permutation. P-values were then obtained from the cumulative distribution function of the fitted Gamma distribution.

### 5.9 Annotation

#### 5.9.1 Locus definition

A locus was considered significant in the discovery cohort if its lead SNP reached the conventional genome-wide significance threshold of *p <* 5 × 10*^−^*^8^, which accounts for the effective number of independent SNPs tested in European-ancestry populations [46]. To further account for the 56 cortical regions tested simultaneously, a Bonferroni correction was applied, yielding a more stringent threshold of 5 × 10*^−^*^8^*/*56 ≈ 8 × 10*^−^*^10^. Because cortical regions partially overlap anatomically, association statistics across regions are positively correlated and therefore not independent. As a result, the Bonferroni correction assuming 56 independent tests is conservative, and overestimates the effective multiple testing burden.

For each region encoded by its representation space, independent lead SNPs and genomic loci were defined following the protocol of Watanabe et al. [47] as implemented in the Functional Mapping and Annotation of Genome-Wide Association Studies platform (FUMA), using PLINK v1.9 [48]. PLINK v2 was not used as it does not support the clumping procedure. In a first step, candidate index SNPs were identified at the primary significance threshold (–clump-p1 5e-8, or –clump-p1 8e-10 for the Bonferroni-corrected threshold). SNPs with *p <* 5 × 10*^−^*^6^ were included in clumps if their LD with the index SNP exceeded *r*^2^ = 0.6 within a 1 Mb window (–clump-p2 5e-6, –clump-r2 0.6, –clump-kb 1000). In a second step, independent lead SNPs were identified by re-applying the clumping procedure to the index SNPs retained in the first step, using a more stringent LD threshold of *r*^2^ = 0.1 with otherwise identical parameters. Clumps separated by less than 250 kb were subsequently merged into a single genomic locus, represented by the lead SNP with the lowest p-value. At each step, the LD was calculated on the population of the study with the flag –bfile.

To enable cross-phenotype comparisons between Champollion V1 regional folding phenotypes and other cortical or subcortical traits reported in the literature, loci identified across the 56 regions were collapsed into a non-redundant set by merging any two loci whose genomic windows overlapped. The merged loci are broader and fewer in number than the region-specific loci, and are each represented by the lead SNP with the lowest p-value prior to merging.

#### 5.9.2 Replication criteria

A lead SNP identified in the discovery cohort was considered replicated if its association with the representation of the same sulcal region in the replication cohort reached nominal significance (*p <* 0.05), following the criterion used in Shadrin at al. [25].

Replication power is primarily constrained by the difference in sample size. For a given effect, the z-score attenuates proportionally to the square root of the sample size ratio: 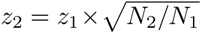. Under this approximation alone, replicating at the nominal threshold *α* = 0.05 in *N*_2_ = 9 950 ABCD participants using associations discovered in *N*_1_ = 35 940 UKB participants requires a minimum discovery z-score of *z*_1_ ≥ 3.72 (i.e. p ≤ 2 × 10*^−^*^4^). However, cross-cohort differences in ancestry, acquisition protocol, and genotyping arrays further attenuate the transferable information beyond what sample size alone would predict. To quantify this additional attenuation, we regressed the z-scores of variants passing the Bonferroni-corrected threshold (*p <* 8 × 10*^−^*^10^) in UKB against their corresponding z-scores in ABCD yields a slope of *z*_2*eff*_ */z*_1*eff*_ = 0.37, corresponding to an effective sample size *N*_2eff_ = (*z*_2*eff*_ */z*_1*eff*_)^2^ × *N*_1_ = 4 884, approximately half the actual ABCD sample size. Under these empirically calibrated conditions, replication at *α* = 0.05 requires a discovery z-score *z*_1_ ≥ 5.32 (*p <* 1 × 10*^−^*^7^). Replication analyses were therefore conducted in the full ABCD sample of 9,950 participants (mean age 9.9 ± 0.6 years, mixed ancestries). Restricting replication to the 5,269 European ancestry ABCD participants would reduce power further, requiring a discovery z-score of *z*_1_ *>* 7.3 (*p <* 2.8*e* 13), a threshold met by only a small number of loci, and was therefore not pursued.

#### 5.9.3 Inter-hemispheric overlap of associated loci

To quantify the overlap between the sets of loci identified independently in the left and right hemispheres for each sulcal region, a Dice Similarity Coefficient (DSC) was computed for each pair of regions. For two sets of genomic loci A and B, each defined by their chromosomal windows [start, end], the DSC is defined as: 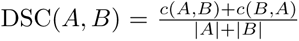 where *c*(*A, B*) counts the number of loci in A that overlap at least one locus in B, with overlap defined as any non-empty intersection of chromosomal windows. This coefficient ranges from 0 (no overlap) to 1. Because loci are defined using LD-based clumping and merging, Dice overlap reflects both shared associations and underlying LD structure.

The DSC was computed for all 56 × 55*/*2 = 1 540 pairs of regions using the post-gwas tools, an in-house Python 3.10 library, applied to the GenomicRiskLoci.txt files at the *p <* 5 × 10*^−^*^8^ threshold. The 28 inter-hemispheric pairs, each corresponding to the left and right instances of the same sulcal region, were extracted from the full 56 × 56 DSC matrix and compared to the distribution of DSC values across all pairs of anatomically distinct regions.

#### 5.9.4 Gene-level enrichment analysis

Gene-level association statistics were computed using MAGMA v1.10 [49]. SNPs were mapped to genes using a 20 kb window upstream and downstream of each gene body (–annotate window=20,20). For UKB discovery cohort, linkage disequilibrium was estimated using the 1000 Genomes Phase 3 European reference panel (g1000_eur), and gene coordinates were defined according to GRCh37, using the Ensembl v102 gene annotation file provided by FUMA (ENSGv102.coding.genes.txt). For ABCD replication cohort, LD was estimated directly from the cohort genotype files (–bfile), and gene positions were defined according to GRCh38.p14 using an annotation file exported from Ensembl BioMart. MAGMA yields a magma.genes.raw file for each summary statistics, containing the gene-level annotation. This gene-level output (magma.genes.raw) was subsequently used as input for the gene-set enrichment, BrainSpan expression enrichment, and single cell enrichment analyses described below.

#### 5.9.5 Gene-set enrichment analysis

Gene-set enrichment was assessed for each region independently against the MSigDB gene set collection [50] (MSigDB_20231Hs_MAGMA.txt, 17,009 gene sets), using the gene-level output (magma.genes.raw; see previous section) as input via –gene-results.

Manhattan plots, QQ plots, and Miami plots were generated using post-gwas tools.

#### 5.9.6 Gene expression enrichment analysis - BrainSpan

To identify developmental time windows during which genes associated with cortical folding representations. are preferentially expressed in the human brain, a gene-property analysis was conducted using MAGMA v1.10 [49] and expression data from the BrainSpan Atlas of the Developing Human Brain [51]. Gene expression covariates were derived from the BrainSpan atlas, with an average log_2_ transformed RPKM expression values for each gene across 29 age points spanning early fetal development (8 post-conception weeks, pcw) to late adulthood (40 years). The gene-property analysis was performed using the –gene-covar option of MAGMA, conditioning on the average expression across all time points (–model direction-covar=greater condition-hide=Average), so that the test evaluates whether genes with higher association statistics are preferentially expressed at each specific developmental stage relative to their overall expression profile. It relies on the gene-level annotation (–gene-results magma.genes.raw). This analysis was conducted independently for each of the 56 sulcal regions.

#### 5.9.7 Meta-analysis across sulcal regions

Since all regions are part of a single whole, and to make the results easier to interpret, results were aggregated using a method adjusted for the correlation structure that exists among the regions.

To combine gene-level and gene-set-level enrichments across the 56 sulcal regions, an empirical inter-region correlation matrix **R** was estimated from the gene-level (magma.genes.raw) files, capturing the statistical dependence at least induced by the partial anatomical overlap between regions. For each gene or gene set, regional z-scores were combined into a single global z-score using a correlation-corrected Stouffer statistic [52]: 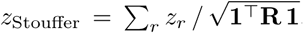, where **1** is the vector of ones of length 56. This denominator accounts for the effective number of independent regions. Global significant threshold were then adjusted for multiple testing using the Bonferroni correction, at *p*_bonf_ *<* 0.05*/*19 264 for genes and *p*_bonf_ *<* 0.05*/*17 009 for gene sets.

To combine BrainSpan results across regions, the one-sided p-values returned by MAGMA for each developmental time point and each region were converted to z-scores as *z* = Φ*^−^*^1^(1 − *p*), where Φ*^−^*^1^ denotes the probit function, preserving the directionality of the enrichment. The same empirical inter-region correlation matrix **R**, estimated from gene-level MAGMA z-scores, was used to apply the correlation-corrected Stouffer aggregation across the 56 regions for each developmental time point. A Bonferroni correction was applied over the 29 time points tested (*p*_bonf_ *<* 0.05*/*29).

#### 5.9.8 Single cell enrichment analysis

To identify the brain cell types in which GWAS-associated genes are preferentially expressed, we performed cell-type enrichment analysis using the MAGMA gene-property framework [49]. This approach tests, for each cell type, whether genes significantly associated with a given phenotype at the gene level tend to be specifically expressed in that cell type, using a linear regression model. Expression specificity was used as a continuous gene-level covariate, with the average expression across all cell types included as a conditioning variable (condition-hide=Average) to control for overall transcriptional activity and avoid confounding by broadly expressed genes. The test was restricted to the positive direction (direction=greater). It relies on the gene-level annotation (–gene-results magma.genes.raw).

Cell-type expression profiles were obtained from the pre-processed scRNA-seq datasets provided by FUMA [53], derived from the human fetal cortex atlas of Bhaduri *et al.* 2021. This atlas characterises transcriptomic diversity across six cortical regions — the Primary Visual cortex, the Temporal cortex, the Primary Somatosensory cortex, the Primary Motor cortex, the Parietal cortex, and the Prefrontal cortex — at developmental time points spanning 14 to 25 post-conception weeks (PCW). This window encompasses key stages of human cortical development, including active neurogenesis, radial migration, cortical layer formation, and the onset of gliogenesis, making it particularly relevant for traits derived from structural brain MRI.

For each phenotype, MAGMA gene-property analysis was performed independently for the corresponding cortical region and all developmental time points. Since multiple cell types were tested within each analysis, p-values were adjusted using the Benjamini-Hochberg procedure to control the false discovery rate (FDR) across all cell types and developmental stage within each region. Corrected enrichment significance was subsequently represented as log_10_(FDR), allowing direct comparison of the temporal dynamics of cell-type enrichment across fetal cortical development.

#### 5.9.9 Comparison of discovered loci with the GWAS Catalog

To evaluate the overlap between the loci identified in this study and previously reported genetic associations, the GWAS Catalog REST API [54] was queried in March 2026 for each lead SNP identified at the genome-wide significance threshold of *p <* 5 × 10*^−^*^8^. For each locus, the rsID lead SNP with the lowest p-value was used as the primary query; if no matching entry was found in the GWAS Catalog, alternative lead SNPs from the same locus were queried in ascending order of pvalue. A locus was considered to overlap a previously reported association if at least one published study in the GWAS Catalog reported an association with the same variant at a conventional genome-wide significance threshold. Studies were then ranked by the number of lead SNPs shared with our discovery set in descending order, in order to identify the phenotypes and cohorts with the greatest genetic overlap with the sulcal folding morphology captured by Champollion V1.

#### 5.9.10 Genetic correlations with psychiatric and neurological disorders

Genetic correlations between the representation spaces of Champollion V1 and a panel of seven psychiatric and neurological disorders were estimated using LD Score Regression (LDSC) [55], with both as LD reference and regression weights (–ref-ld-chr and –w-ld-chr, eur_w_ld_chr), after summary statistics were harmonised using the munge_sumstats.py script. The disorders included were: schizophrenia, bipolar disorder, attention-deficit/hyperactivity disorder (ADHD), autism spectrum disorder (ASD), depression, neuroticism, and post-traumatic stress disorder (PTSD). Summary statistics for each disorder were obtained from the following sources: schizophrenia [56], bipolar disorder [57], ADHD [58], ASD [59], PTSD [60], and neuroticism and depression [61].

For each sulcal region and each disorder, genetic correlations were estimated independently for the first three principal components of the regional representation space using the corresponding univariate GWAS summary statistics. To combine evidence across the three dimensions into a single global test, we computed a Mahalanobis-type *χ*^2^ statistic that explicitly accounts for the genetic correlation structure among principal components. Specifically, let **z** = (*z*_1_*, z*_2_*, z*_3_)*^⊤^* denote the vector of LDSC *z*-scores for the genetic correlation between each dimension and the disorder of interest, and let **Σ** denote the 3 × 3 matrix of pairwise genetic correlations among the three principal components, estimated from the same LDSC framework. The omnibus statistic is then: *χ*^2^ = **z***^⊤^***Σ***^−^*^1^**z** which follows an approximate *χ*^2^ distribution with three degrees of freedom under the global null hypothesis that the disorder is genetically uncorrelated with all three dimensions. In cases where the estimated **Σ** matrix was not positive semi-definite (likely attributable to sampling noise in LDSC pairwise estimates for low-heritability regions), the region is removed. Affected regions are in white in Extended Data Fig. 13.

This global test evaluates whether any linear combination of the first three principal components of a given regional representation space is genetically correlated with the disorder of interest. A sulcal region was considered significantly correlated with a disorder if the resulting *χ*^2^ p-value survived Bonferroni correction for the 56 × 7 = 392 tests performed (*p*_bonf_ *<* 0.05*/*392 = 1.28 × 10*^−^*^4^).

The present genetic correlation analyses were restricted to the first three principal components of each regional representation space. This choice was made because 3 PCs already captured a large part of the genetic variance (see Extended Data Fig. 14). Also, a sensitivity analysis was conducted to see the influence of more PCs (see Extended Data Fig. 13), confirming the correlation found on only three PCs, when using 4 PCs, 5 PCs, and 6 PCs. The only issue is the stability of the genetic correlation matrix **Σ**, which tends not to be positive semi-definite, which creates then aberrant results for regions with poorly estimated genetic correlation matrices, because of a very low heritability.

## 6 Code availability

Here you will find repositories for:

BrainVISA: https://brainvisa.info/web/

Cortical tiles: https://github.com/neurospin/cortical_tiles/releases/tag/canonical_25

Champollion V1: https://huggingface.co/neurospin/Champollion_V1, https://github.com/neurospin/champollion/releases/tag/champollion_v1_canonical_25

Decoder: https://github.com/neurospin-projects/2025_adufournet_champollion_decoder/tree/v0.0.0

MOSTest: https://github.com/precimed/mostest

Post-GWAS tools: https://github.com/neurospin/postgwas-tools/releases/tag/v1.0.0

PLINK 1.9: https://www.cog-genomics.org/plink/1.9/

PLINK 2.0: https://www.cog-genomics.org/plink/2.0/

MAGMA v1.10: https://cncr.nl/research/magma/

FUMA 2.0.0: https://fuma.ctglab.nl/

LDSC v2.0.1: https://github.com/bulik/LDSC

GWAS Catalog REST API: https://www.ebi.ac.uk/gwas/rest/docs/api

## 7 Data availability

This research was conducted using the UK Biobank resource under application number 64984.

Data used in the preparation of this article were obtained from the Adolescent Brain Cognitive Development™ (ABCD) Study https://abcdstudy.org/, held in the NIH Brain Development Co-horts Data Sharing Platform https://www.nbdc-datahub.org/. This is a multisite, longitudinal study designed to recruit more than 10,000 children aged 9-10 and follow them over 10 years into early adulthood.

The ABCD Study is supported by the National Institutes of Health and additional federal partners under award numbers: U01DA041048, U01DA050989, U01DA051016, U01DA041022, U01DA051018, U01DA051037, U01DA050987, U01DA041174, U01DA041106, U01DA041117, U01DA041028, U01DA041134, U01DA050988, U01DA051039, U01DA041156, U01DA041025, U01DA041120, U01DA051038, U01DA041148, U01DA041093, U01DA041089, U24DA041123, U24DA041147.

A full list of supporters is available at Federal Partners - ABCD Study https://abcdstudy.org/federal-partners.html.

ABCD Consortium investigators designed and implemented the study and/or provided data but did not necessarily participate in the analysis or writing of this report. This manuscript reflects the views of the authors and may not reflect the opinions or views of the NIH or ABCD Consortium investigators.

BrainSpan: https://www.brainspan.org/

Summary statistics (change the GCST according to the file you want to download, see the gcst_list.xlsx): https://www.ebi.ac.uk/gwas/studies/GCST90841311

## 8 Author contributions

A. Dufournet, V. Frouin and J.-F. Mangin conceived the research idea. D. Rivière and J. Chavas provided advice on study design. J. Laval, J. Chavas and A. Dufournet pretrained the models and generated the self-supervised representations of cortical folding. C. Fisher and D. Rivière provided the classical sulcal morphometric traits. A. Dufournet and V. Frouin performed the mGWAS analysis, the LDSC analysis, the MAGMA analyses, and the single-cell analysis. A. Dufournet wrote the manuscript. All authors read and approved the manuscript.

## 9 Acknowledgments

This project has been funded by ANR via FOLDDICO (ANR-20-CHIA-0027), BHT (ANR-22-PESN-0012) and StratifyAging (ANR-22-PESN-0010), RHU-PsyCARE (ANR-18-RHUS-0014), the Audace program of the CEA (Cortical folding), by European Union’s Horizon 2020 for R-LiNK (H2020-SC1-2017, 754907), and EBRAINS2 (HORIZON-INFRA-2022-SERV-B-01). This project was provided with computing HPC and storage resources by GENCI at TGCC thanks to the grant 2024-A0170313800 on the supercomputer Joliot Curie’s ROME partition. This project was provided with computer and storage resources by GENCI at IDRIS thanks to the grant AD010316018 on the supercomputer Jean Zay’s V100 partition.

Claude AI helped with the drafting of this article, to improve readability, grammar, and formatting.

We would like to thank Clément Langlet, Saeb Tounsi, Cristobal Mendoza, Abdelghani Baroud, Timothée Sanchez, Noah Tournier, Ivan Bautista, Ege Kibrislioglu, Lilian Marsal, Mathys Bodelet, Barthélémy Drabczuk, Théo Delmaire, Antoine Didier, Racim Menasria, and Pauline Amrouche for their feedback, which helped improve this paper.

