## Extended Data+ for "Self-supervised representations reveal the genetic architecture of human cortical folding"

### 11 Extended Data

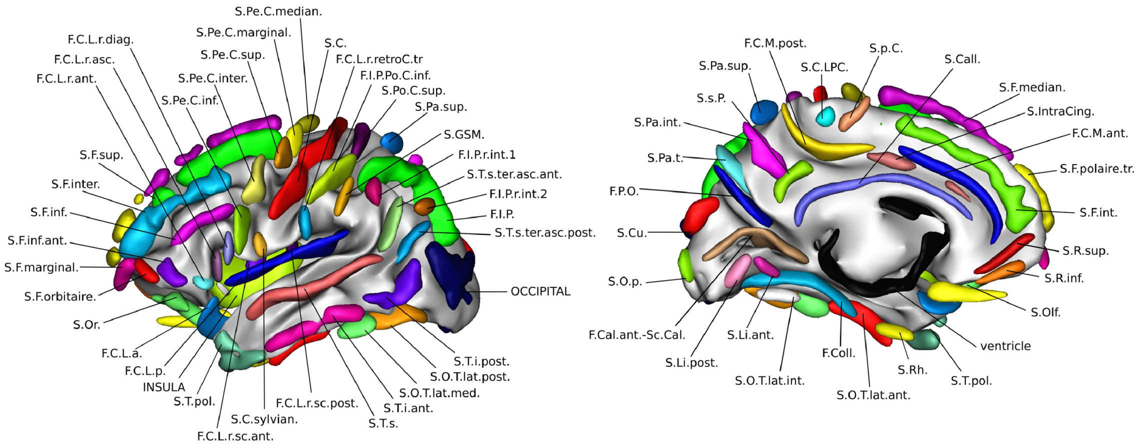


Figure 7: Atlas representing the location of the sulcal labels used in Champollion V1 for the construction of the regions, mapped on a 3D representation of a left hemisphere. Note that right hemisphere atlas is bilaterally symmetric.

#### 11.1 Comparison of MOSTest and MinP

As a validation analysis, the MinP approach was applied in parallel to the 56 cortical regions and identified 299 and 161 independent loci after merging the results at the respective thresholds of $5 \times 10^{-8}$ and $8 \times 10^{-10}$. More than 90% of these loci overlapped with those identified using MOSTest, supporting the robustness of the associations across multivariate testing strategies.

#### 11.2 Empirical false discovery rate

To empirically estimate the false discovery rate (FDR) at the SNP level without accounting for linkage disequilibrium (LD), we leveraged the permutation framework implemented in MOSTest [23]. Briefly, for each SNP in each region, a single permuted genotype vector is generated, then the association test across the 32 dimensions is computed, to yield a null distribution of test statistics while maintaining the phenotypic correlation structure. Across $8\,109\,955 \times 56$ tests, the empirical false discovery rate at the $5 \times 10^{-8}$ threshold was $22/333\,048 = 6.6 \times 10^{-5}$, as 22 SNPs exceeded the threshold on average across the 56 cortical regions under permutation, compared with 333,048 significant SNPs across all region prior to LD clumping. This rate is two orders of magnitude below the nominal 5% level, confirming control of type I error. At the more stringent threshold of $8 \times 10^{-10}$, no false positives were observed across the $8\,109\,955 \times 56$ SNP permutations performed. Critically, this permutation scheme, one permuted genotype vector per SNP, reused across all phenotypes, differs from traditional phenotype label permutations but is theoretically justified for the MOSTest [23].

The genomic inflation factor ($\lambda_{GC}$), estimated for each cortical region in the UK Biobank cohort, ranged from 1.17 to 1.43 for the MOSTest results. To distinguish inflation arising from technical bias versus true polygenic trait, LD Score Regression (LDSC) intercepts were computed across all dimensions and regions [55]. The resulting intercepts, close to 1.00 (Supplemental Material Fig.2), indicate that most of the observed inflation in $\lambda_{GC}$ is attributable to polygenicity rather than confounding artifacts, supporting appropriate control of type I error beyond the permutation-based analysis described above.

#### 11.3 Self-supervised learned representation VS sulcus width

To further validate the specificity of Champollion V1 representations, we conducted a comparison using sulcus opening (sulcus width), a sulcal parameter known to be phenotypically uncorrelated with Champollion V1 [8]. Unlike the morphometric parameters considered above, sulcal opening is regarded as a marker of brain biological aging rather than neurodevelopment: sulci progressively widen with age as a result of cortical atrophy, and when residualized for age and sex, inter-individual variation in sulcal width reflects the degree to which a brain appears older or younger than expected. This serves as a negative control: if the genetic enrichment observed above reflects biological overlap, orthogonal phenotypes should yield largely orthogonal associations. Consistent with this prediction, sulcus opening shows a large proportion (70 out of 111 independent loci) of its associations orthogonal to those of Champollion V1 (Extended Data Fig. 8). The notable exception is chromosome 17, where shared associations persist, likely reflecting the well-documented broad pleiotropy of this region across brain phenotypes.

#### 11.4 Comparison with the GWAS catalog

To assess the novelty and biological relevance of the identified loci, we queried the GWAS Catalog API for each lead SNP, retrieving previously reported associations from published studies. The five studies with the greatest overlap with our discovery set are all imaging genetics studies. The largest overlap is with [23], which reported associations with brain morphology attributes ($n = 26\ 502$ UKB white British ancestry individuals, GCST010703), sharing 164 lead SNPs with our results, followed by the same study for cortical surface area specifically (GCST010701), with 125 shared lead SNPs. Cortical thickness [33] ($n = 33\ 748$ UKB white British ancestry individuals, GCST90095131) and a broad neuroimaging measurement study [62] ($n = 34\ 029$ UKB white British ancestry individuals, GCST90319488) follow with 79 and 70 shared lead SNPs respectively. Notably, the volume of Brodmann Area 3b in the right hemisphere [63] ($n = 21\ 281$ UKB white British ancestry individuals, GCST90002852) shares 25 lead SNPs with our results. To our knowledge, 188 independent loci have not been reported in the GWAS Catalog as of March 2026, suggesting novelty relative to published cortical imaging genetics studies.

A more detailed comparison with the results of [23], obtained using conventional cortical phenotypes (cortical thickness, surface area, and subcortical volumes) in 26,502 participants, further supports the biological relevance of the associations identified here. At the genome-wide significance threshold ($5 \times 10^{-8}$), 70% of the 344 loci reported in that study overlap with those identified using Champollion V1 representations (Extended Data Fig. 9a, 9b). A scatter plot of $-\log_{10}(p)$ values nevertheless indicates that the two phenotypic frameworks capture partly distinct associations (Extended Data Fig. 9c).

#### 11.5 Strongest loci

Among all loci identified across the 56 regions, the five most significant LD-independent associations are located in genes known to play roles in cortical and neuronal development and span diverse anatomical regions. The strongest association was observed at chromosome 15q14, represented by rs2033939 ($p = 6 \times 10^{-322}$ in SC-SPeC left), a locus previously associated with classical sulcal morphometric traits [24]. The second strongest association was located at chromosome 14q23.1, represented by rs74826997 ($p = 6 \times 10^{-131}$ in FPO-SCu-ScCal left), within the *DAAM1* gene, which is involved in neurite outgrowth and cell motility [64] and has been linked to multiple structural imaging traits. The third locus, rs13107325 ($p = 6 \times 10^{-91}$ in FCLp-subsc-FCLa-INSULA right), corresponds to *SLC39A8*, a highly pleiotropic variant previously associated with a wide range of psychiatric and cognitive phenotypes [65]. The fourth association, rs12146713 ($p = 6 \times 10^{-90}$ in SCall right), is linked to *NUAK1*, a gene recently associated with glymphatic function and neurodevelopmental disorders [66]. Finally, rs2009778 ($p = 2 \times 10^{-84}$ in SFinf-BROCA-SPeCinf left), located on chromosome 2p14, has previously been reported in the anterior cingulate cortex [22].

Importantly, the observed associations were highly stable with respect to population structure adjustment. Using either 10 or 20 genetic principal components as covariates yielded highly consistent results across all cortical regions, indicating that the findings are unlikely to be driven by residual stratification effects.

#### 11.6 Genetic Dice similarity across regions

The Dice similarity coefficient computed on independent loci reaching genome-wide significance ($5 \times 10^{-8}$) at the regional level reveals a strong inter-hemispheric symmetry. For a given region, the overlap (defined as intersecting genomic loci; see Methods) between loci identified in the left and right hemispheres is $0.62 \pm 0.06$ on average, which is 2.2 times greater than the mean overlap observed between distinct regions ($0.28 \pm 0.13$).

This result indicates that largely overlapping sets of genetic variants influence cortical folding morphology in a broadly symmetric manner across hemispheres. Symmetry is highest for SOr (0.71) and lowest for SFint-SR (0.44), the latter also being the least heritable region, consistent with shared genetic influences across hemispheres, although part of this effect may reflect differences in statistical power across regions.

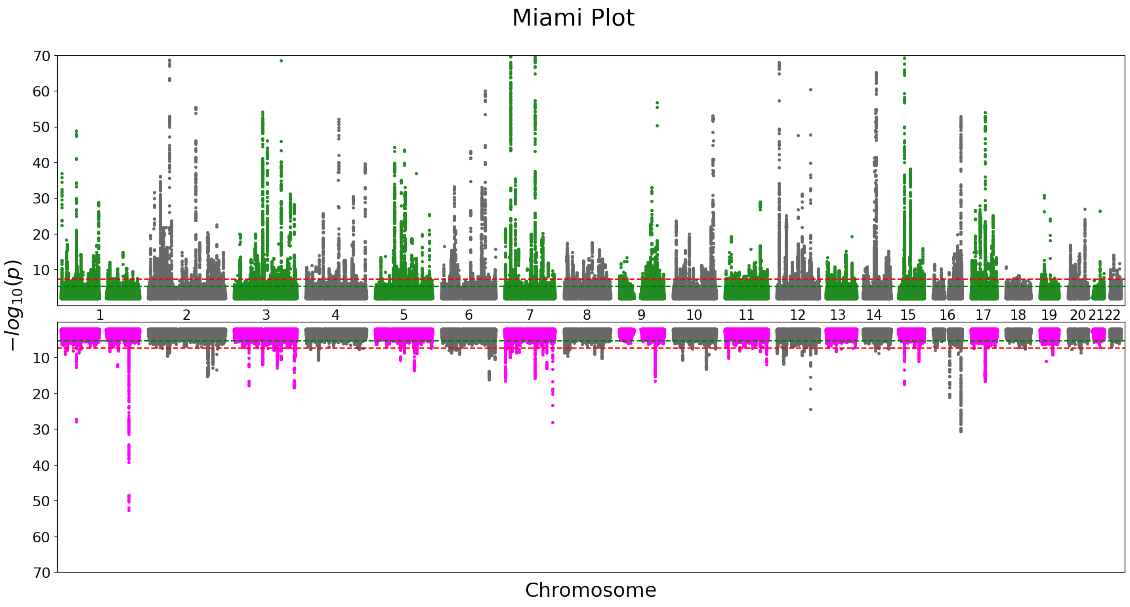


(a)

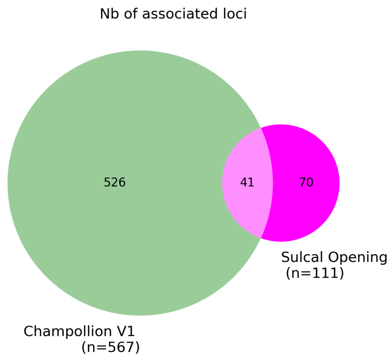


(b)

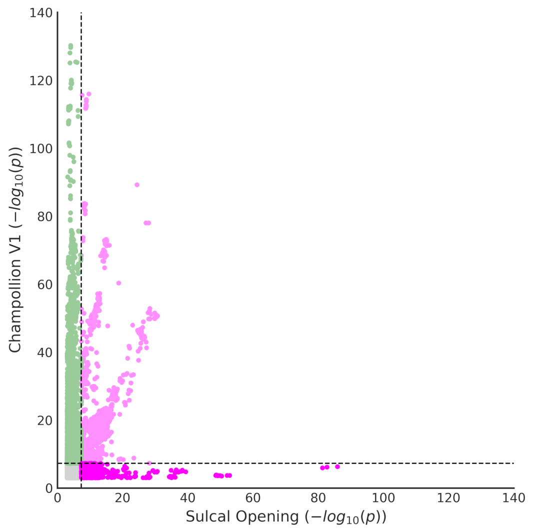


(c)

Figure 8: Champollion V1 representations are orthogonal to sulcal opening. **(a)** Miami plot comparing the strongest associations (using MOSTest) for all regions encoded by Champollion V1 on the 35,940 UKB white British ancestry subjects (green) to the associations related to sulcal opening (magenta). The conventional threshold of $5 \times 10^{-8}$ is indicated with the red line. Note that the window is clipped for the $-\log_{10}$(p-value) above 70. **(b)** Overlap (pink) of significant loci between Champollion V1 (green) and sulcal opening (magenta). Overlap is considered to exist if there is at least one significant SNP within both loci. Numbers indicate the count of independent merged loci unique to each method or shared between them. **(c)** Scatterplot of SNP-based $-\log_{10}(p)$, with y-axis for Champollion V1 and x-axis for sulcal opening. The green points correspond to SNPs whose association appears only in the Champollion V1 representations. The magenta points correspond to SNPs whose association appears only in the sulcal opening. The conventional threshold of $5 \times 10^{-8}$ is indicated with the black dotted line. Note that $-\log_{10}(p)$ are clipped at 140 on both axes.

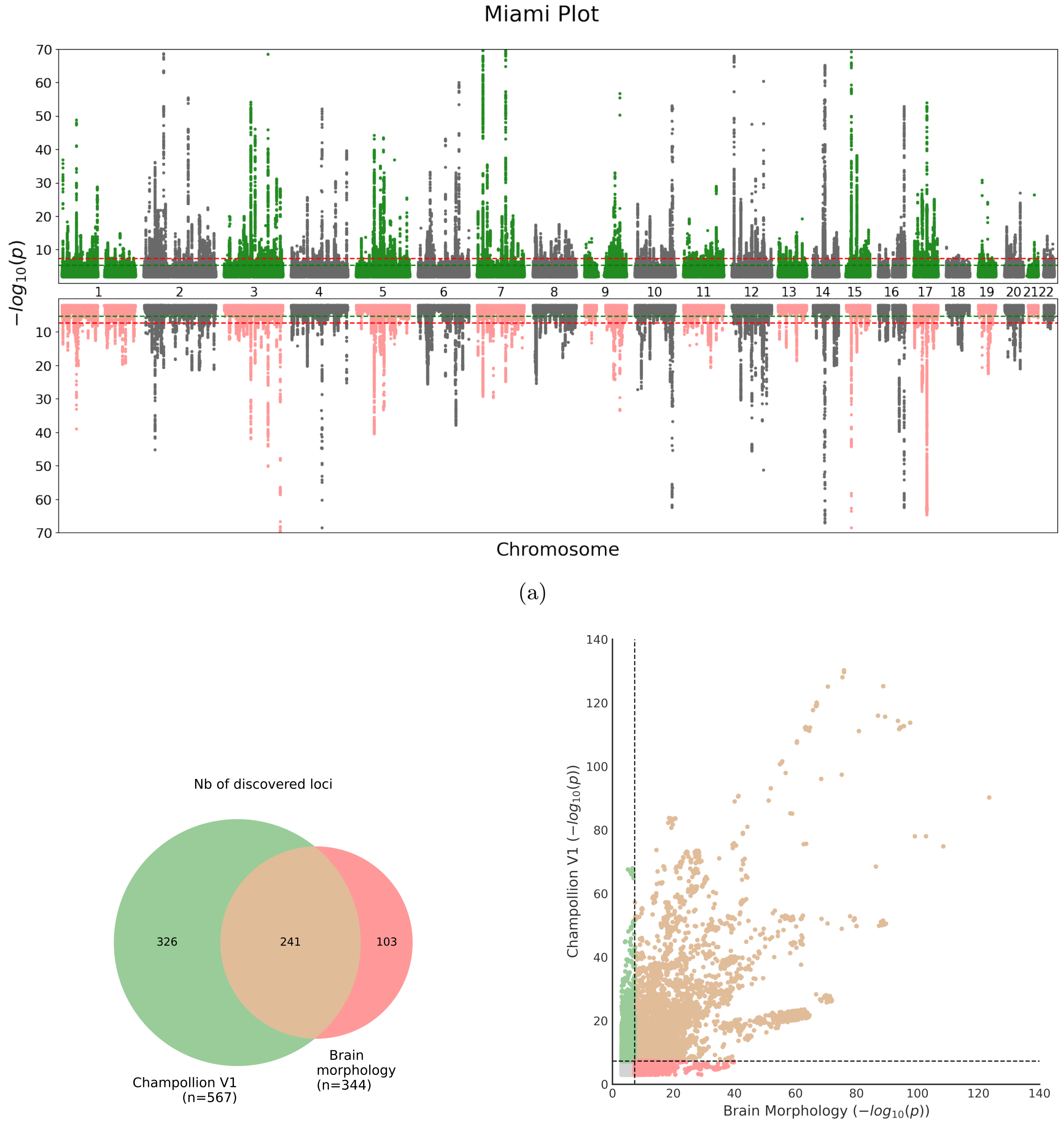


(a) (b) (c)

Figure 9: Comparison of genetic associations between Champollion V1 and brain morphology (subcortical volumes, cortical thickness and surface area) from [23]. **(a)** Miami plot comparing the strongest associations (using MOSTest) for all regions encoded by Champollion V1 on the 35,940 UKB white British ancestry subjects (green) to the associations from the brain morphology on the 26,502 white British ancestry subjects [23] (pink). Note that the window is clipped for the $-\log_{10}$(p-value) above 70. **(b)** Overlap of significant loci between Champollion V1 (green) and brain morphology (subcortical volumes, cortical thickness and surface area) (pink). Overlap (light brown) is considered to exist if there is at least one significant SNP ($p < 5 \times 10^{-8}$) within both loci. **(c)** Scatterplot of SNP-based $-\log_{10}(p)$, with y-axis for Champollion V1 and x-axis for brain morphology. The conventional threshold of $5 \times 10^{-8}$ is indicated with the black dotted line. Note that $-\log_{10}(p)$ are clipped at 140.

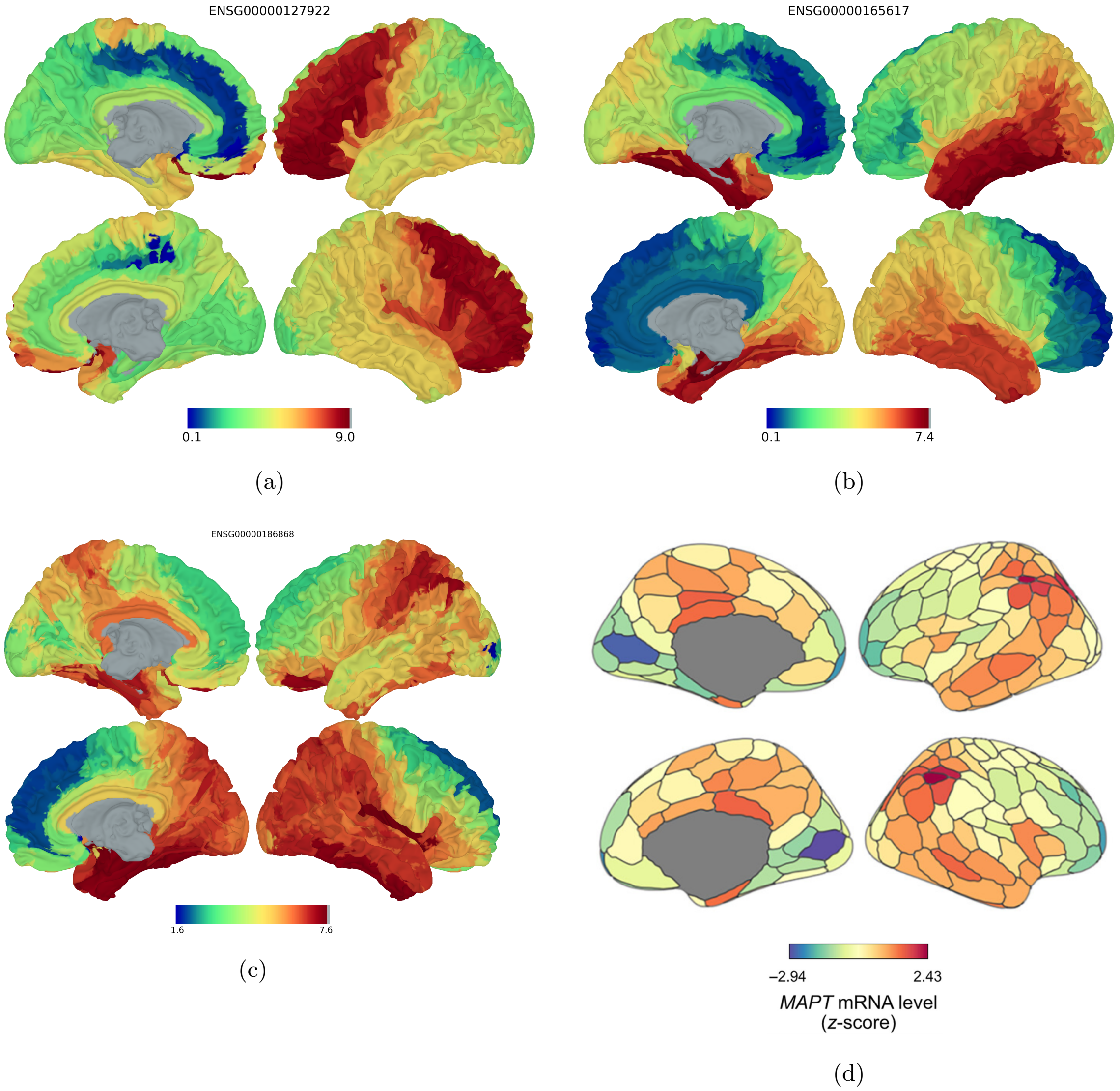


Figure 10: Cortex-wide `MAGMA` gene association maps for *SEM1* (panel **a**), for *DACT1* (panel **b**), and *MAPT* (panel **c**). For each gene, `MAGMA` z-scores quantifying gene-level association with cortical folding morphology (encoded in Champollion V1 representations) are projected onto the cortical surface (MNI ICBM 152 white mesh, four views). Higher z-scores (warmer colours) indicate stronger association. Note that the colour scale differs between panels; direct visual comparison of magnitudes across panels is not appropriate. *SEM1* - ENSG00000127922 shows frontal enrichment and *DACT1* - ENSG00000165617 shows temporal enrichment. (panel **d**) Average MAPT gene expression on the Schaefer atlas, derived from mRNA profiling of 6 healthy human brains in the Allen Human Brain Atlas (AHBA), from the study [28].

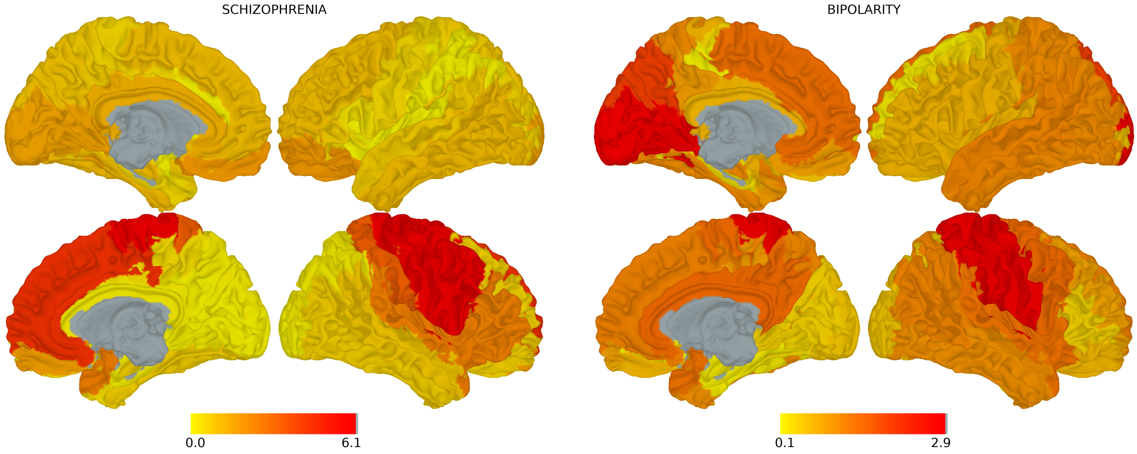


(a) (b)

Figure 11: Genetic correlation maps between cortical folding and psychiatric disorders ($-\log_{10}$(p-value)), represented on the MNI ICBM 152 white mesh (left lateral, left medial, right lateral, and right medial views). Note that the results shown are prior to Bonferroni correction and should not be interpreted as significant for Bipolar disorder. **(a)** Schizophrenia. **(b)** Bipolar disorder.

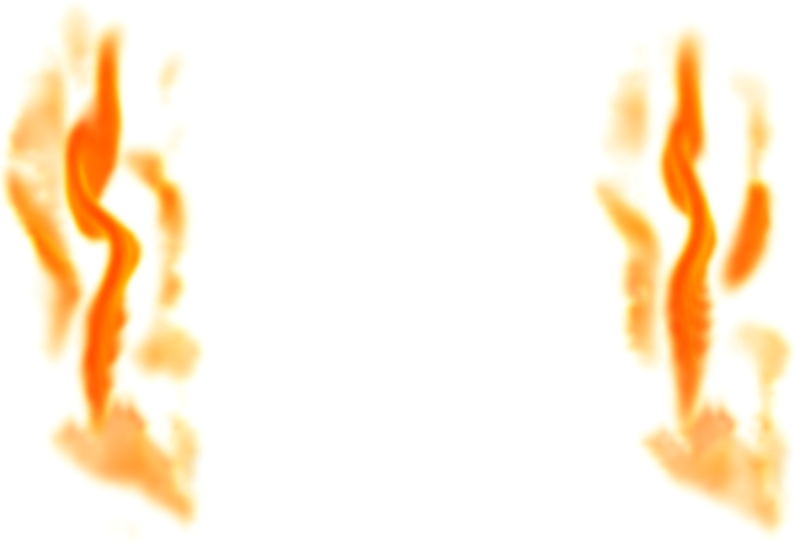

(a) (b)

Figure 12: Examination of the representation space encoding the SC-sylv right region with the reconstruction of the second principal component axis at the **(a)** left end of the principal axis (positive correlation with schizophrenia) and **(b)** right end of the principal axis (negative correlation with schizophrenia)for a given subject.

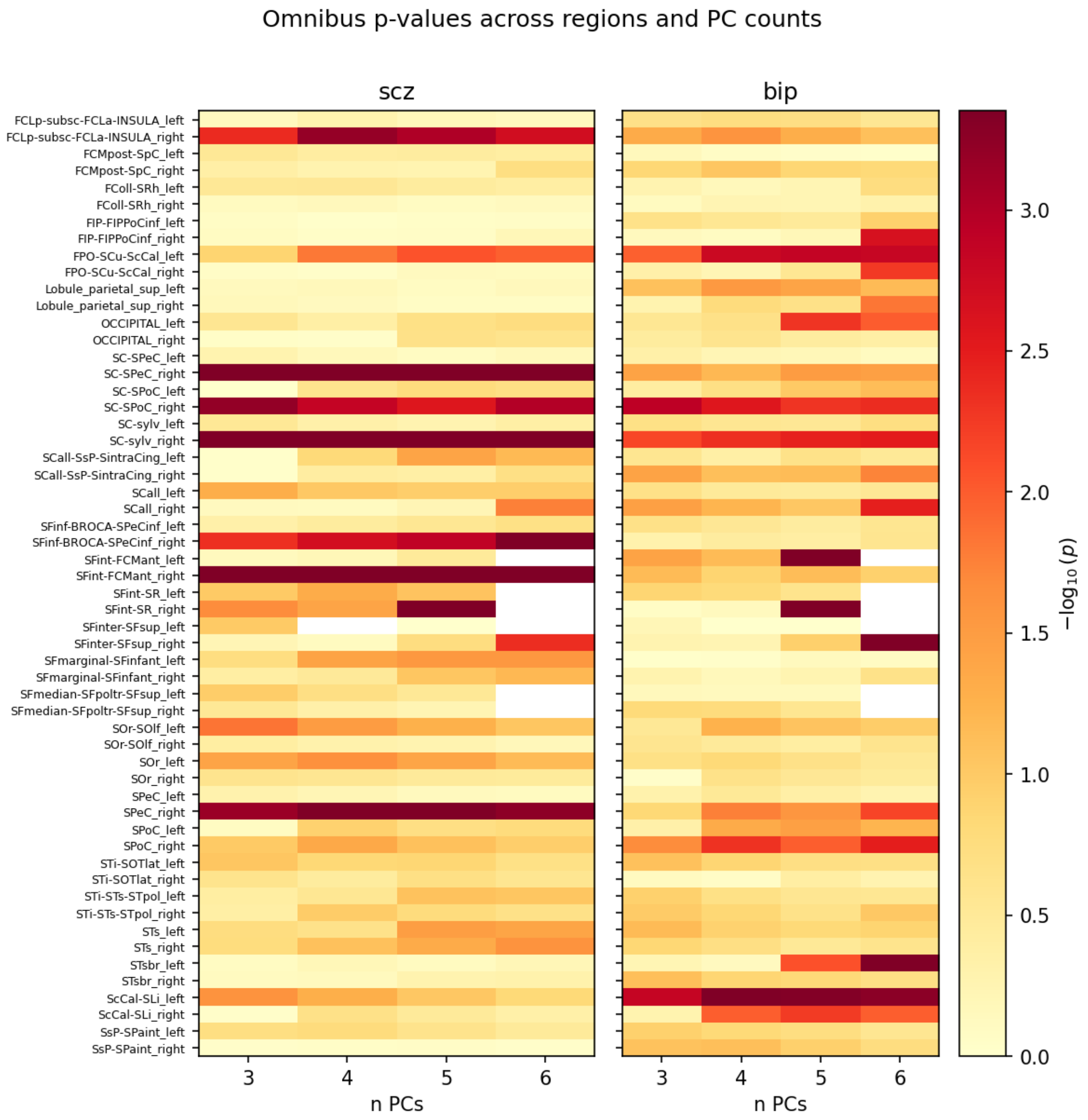


Figure 13: Sensitivity heatmap representing the p-values across region with a significant genetic correlation with schizophrenia and bipolar disorder, according to the number of principal components used for the omnibus test. Note: SFint-FCMant left, SFint-SR right, and SFinter-SFsup regions had ill-conditioned $\mathbf{\Sigma}$ matrices (likely attributable to low heritability), which may produce inflated or unstable genetic correlation estimates.

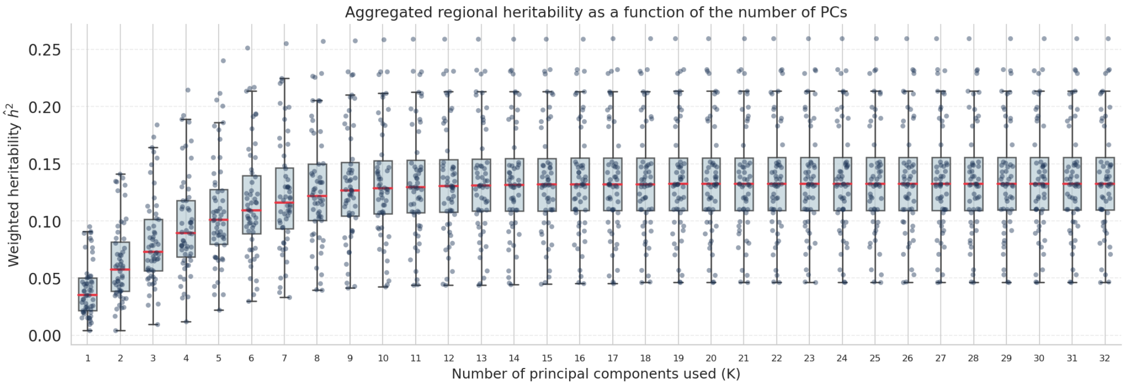


Figure 14: Estimation of aggregate heritability based on SNPs as a function of the number of retained principal components. Each point represents the aggregate heritability of a region. With this method, components that explain very little variance in the original space are also of little importance for heritability.

| Region | SNP-based $h^2$ | SE |
|---|---|---|
| SCall-SsP-SintraCing left | 0.259 | 0.009 |
| SCall left | 0.233 | 0.008 |
| FPO-SCu-ScCal right | 0.231 | 0.009 |
| FPO-SCu-ScCal left | 0.229 | 0.009 |
| SCall-SsP-SintraCing right | 0.213 | 0.008 |
| ScCal-SLi right | 0.213 | 0.009 |
| SCall right | 0.211 | 0.008 |
| FCLp-subsc-FCLa-INSULA left | 0.200 | 0.008 |
| ScCal-SLi left | 0.198 | 0.007 |
| FColl-SRh left | 0.192 | 0.007 |
| FCLp-subsc-FCLa-INSULA right | 0.190 | 0.007 |
| FColl-SRh right | 0.188 | 0.006 |
| STi-SOTlat right | 0.183 | 0.006 |
| STi-SOTlat left | 0.167 | 0.006 |
| SC-sylv right | 0.152 | 0.008 |
| STi-STs-STpol left | 0.151 | 0.006 |
| SC-SPoC right | 0.149 | 0.007 |
| SC-sylv left | 0.149 | 0.007 |
| FIP-FIPPoCinf left | 0.148 | 0.006 |
| Lobule parietal sup left | 0.147 | 0.006 |
| Lobule parietal sup right | 0.143 | 0.006 |
| SC-SPoC left | 0.143 | 0.007 |
| SsP-SPaint left | 0.139 | 0.007 |
| SsP-SPaint right | 0.139 | 0.006 |
| FIP-FIPPoCinf right | 0.137 | 0.006 |
| SOr-SOlf right | 0.137 | 0.006 |
| OCCIPITAL right | 0.133 | 0.006 |
| SOr right | 0.133 | 0.006 |
| STs left | 0.132 | 0.006 |
| SPoC right | 0.132 | 0.006 |
| STi-STs-STpol right | 0.132 | 0.006 |
| SOr-SOlf left | 0.130 | 0.005 |
| OCCIPITAL left | 0.130 | 0.006 |
| STs right | 0.124 | 0.006 |
| SPoC left | 0.121 | 0.006 |
| SC-SPeC left | 0.120 | 0.006 |
| SOr left | 0.120 | 0.005 |
| STsbr left | 0.118 | 0.006 |
| SFinf-BROCA-SPeCinf right | 0.111 | 0.006 |
| FCMpost-SpC right | 0.110 | 0.005 |
| SFinf-BROCA-SPeCinf left | 0.110 | 0.006 |
| SC-SPeC right | 0.110 | 0.006 |
| FCMpost-SpC left | 0.107 | 0.005 |
| STsbr right | 0.100 | 0.005 |
| SPeC right | 0.099 | 0.005 |
| SPeC left | 0.096 | 0.005 |
| SFmarginal-SFinfant left | 0.092 | 0.005 |
| SFmarginal-SFinfant right | 0.089 | 0.005 |
| SFint-FCMant left | 0.083 | 0.005 |
| SFinter-SFsup right | 0.081 | 0.005 |
| SFint-FCMant right | 0.077 | 0.005 |
| SFinter-SFsup left | 0.071 | 0.005 |
| SFint-SR left | 0.057 | 0.005 |
| SFint-SR right | 0.053 | 0.005 |
| SFmedian-SFpoltr-SFsup left | 0.047 | 0.004 |
| SFmedian-SFpoltr-SFsup right | 0.047 | 0.004 |

Table 3: SNP-based $h^2$ aggregated for each regional representation space. See Figure 3b.

| Region | Lead SNPs (discovery) | Lead SNPs (replication) | Replication (%) |
|---|---|---|---|
| FCLp-subsc-FCLa-INSULA left | 48 | 21 | 43.8 |
| FCLp-subsc-FCLa-INSULA right | 52 | 23 | 44.2 |
| FCMpost-SpC left | 15 | 7 | 46.7 |
| FCMpost-SpC right | 15 | 9 | 60.0 |
| FColl-SRh left | 65 | 37 | 56.9 |
| FColl-SRh right | 48 | 30 | 62.5 |
| FIP-FIPPoCinf left | 55 | 39 | 70.9 |
| FIP-FIPPoCinf right | 52 | 29 | 55.8 |
| FPO-SCu-ScCal left | 69 | 36 | 52.2 |
| FPO-SCu-ScCal right | 71 | 42 | 59.2 |
| Lobule parietal sup left | 51 | 30 | 58.8 |
| Lobule parietal sup right | 49 | 19 | 38.8 |
| OCCIPITAL left | 30 | 20 | 66.7 |
| OCCIPITAL right | 39 | 20 | 51.3 |
| SC-SPeC left | 29 | 12 | 41.4 |
| SC-SPeC right | 30 | 13 | 43.3 |
| SC-SPoC left | 39 | 17 | 43.6 |
| SC-SPoC right | 36 | 19 | 52.8 |
| SC-sylv left | 23 | 13 | 56.5 |
| SC-sylv right | 22 | 12 | 54.5 |
| SCall-SsP-SintraCing left | 69 | 44 | 63.8 |
| SCall-SsP-SintraCing right | 55 | 26 | 47.3 |
| SCall left | 47 | 24 | 51.1 |
| SCall right | 48 | 26 | 54.2 |
| SFinf-BROCA-SPeCinf left | 25 | 13 | 52.0 |
| SFinf-BROCA-SPeCinf right | 32 | 18 | 56.2 |
| SFint-FCMant left | 7 | 1 | 14.3 |
| SFint-FCMant right | 11 | 5 | 45.5 |
| SFint-SR left | 4 | 0 | 0.0 |
| SFint-SR right | 7 | 1 | 14.3 |
| SFinter-SFsup left | 18 | 8 | 44.4 |
| SFinter-SFsup right | 28 | 15 | 53.6 |
| SFmarginal-SFinfant left | 17 | 9 | 52.9 |
| SFmarginal-SFinfant right | 22 | 14 | 63.6 |
| SFmedian-SFpoltr-SFsup left | 6 | 0 | 0.0 |
| SFmedian-SFpoltr-SFsup right | 4 | 2 | 50.0 |
| SOr-SOlf left | 56 | 23 | 41.1 |
| SOr-SOlf right | 51 | 28 | 54.9 |
| SOr left | 51 | 25 | 49.0 |
| SOr right | 41 | 22 | 53.7 |
| SPeC left | 24 | 11 | 45.8 |
| SPeC right | 26 | 12 | 46.2 |
| SPoC left | 26 | 9 | 34.6 |
| SPoC right | 27 | 12 | 44.4 |
| STi-SOTlat left | 52 | 30 | 57.7 |
| STi-SOTlat right | 58 | 33 | 56.9 |
| STi-STs-STpol left | 49 | 25 | 51.0 |
| STi-STs-STpol right | 38 | 21 | 55.3 |
| STs left | 32 | 14 | 43.8 |
| STs right | 27 | 8 | 29.6 |
| STsbr left | 39 | 17 | 43.6 |
| STsbr right | 21 | 15 | 71.4 |
| ScCal-SLi left | 91 | 43 | 47.3 |
| ScCal-SLi right | 77 | 42 | 54.5 |
| SsP-SPaint left | 36 | 19 | 52.8 |
| SsP-SPaint right | 33 | 16 | 48.5 |

Table 4: Independent loci identified in UKB (discovery) at $p < 8 \times 10^{-10}$, characterised by their lead SNPs, number of replicated lead SNPs at $p < 0.05$ in ABCD, and replication percentage per region. Only variants common to UKB and ABCD were tested here, which can result in fewer lead SNPs than the total number of genomic loci for a region.
